# WattmaMod enables high-resolution and extensible RNA modification profiling for nanopore direct RNA sequencing

**DOI:** 10.64898/2026.07.02.735990

**Authors:** Bin Yu, Xinghui Sun, Xiao Li, Junhai Qi, Ting Yu, Xin Gao, Renmin Han

## Abstract

Nanopore direct RNA sequencing enables direct profiling of RNA modifications on native transcripts, but accurate multi-modification detection remains limited by non-stationary signals and heterogeneity across chemistries. Here, we develop WattmaMod, a deep learning framework for multi-modification detection from nanopore direct RNA sequencing data. It combines self-supervised pretraining, supervised contrastive fine-tuning, and low-label incremental adaptation to improve representation learning and support efficient extension to low-resource modification types. The framework further incorporates wavelet-guided multi-scale encoding and dynamic cross-attention fusion to model raw signals and event-level features. Results show that WattmaMod achieves robust detection of multiple RNA modifications, including m6A, m5C, m1A, A-to-I, m7G, hm5C, m1Ψ, f5C, ac4C, m5U and Ψ. It also extends efficiently to low-resource modification types with minimal labeled data, generalizes across sequencing chemistries and species, and predicts potential higher-order local organization among distinct RNA modifications. WattmaMod thus provides a scalable framework for high-resolution epitranscriptome profiling and expands RNA modification analysis beyond single-site prediction to coordinated multi-modification characterization.

---

Dynamic RNA modifications regulate RNA processing, stability and translation ^1–6^, and their dysregulation contributes to cancer initiation and progression ^7–9^. However, it remains challenging to systematically profile the distribution and crosstalk of multiple modifications on native RNA molecules ^10–12^. Conventional techniques, including MeRIP-seq ^13^, miCLIP ^14^, m6ACE-Seq ^15^, GLORI ^16^ rely on antibody enrichment or chemical conversion and typically require RNA fragmentation, thereby disrupting long-range transcript information and molecular heterogeneity. Nanopore direct RNA sequencing (DRS) enables real-time, amplification-free sequencing of native RNA ^17^, offering the potential to profile multiple modifications on single molecules ^18–20^. Quantitative deconvolution of distinct modifications from raw current signals remains a bottleneck for broader nanopore DRS applications ^21,22^.

Computational methods for nanopore DRS modification detection largely fall into two paradigms. Comparative statistical tools, including xPore ^23^, DRUMMER ^24^ and Nanocompore ^25^, identify candidate sites by comparing signal distributions between conditions, but their performance depends on well-matched designs and high coverage, limiting single-sample analyses and robustness to batch effects. In parallel, supervised deep-learning models such as m6Anet ^26^, DENA ^27^ and SingleMod ^28^ enable single-nucleotide prediction for m6A, motivating extensions toward multi-modification detection. However, existing multi-modification models, such as TandemMod ^29^, CHEUI ^30^, DirectRM ^31^, and the recently proposed ORCA ^32^, still face practical challenges in cross-chemistry generalization and in high-precision extension to novel modification types. Notably, although ORCA supports multi-modification detection, further improvement is still needed in its detection accuracy and extensibility to additional modification types. These limitations underscore the need for a unified framework supporting multiple chemistries, extensible to new modification types and robust across domains.

To detect RNA modifications, deep learning models must move beyond designed event features to extract robust multiscale representations directly from raw signals, thereby decoupling subtle modification-induced perturbations from background noise ^33–36^. The architecture must remain extensible across nanopore chemistries and invariant to sequence context, enabling efficient adaptation to new modification types or experimental settings ^37^. While RNA004 delivers superior signal quality, the paucity of curated, labeled training data for this new iteration poses a substantial bottleneck to effective model optimization. Furthermore, the framework should generate multi-granular quantitative outputs at both single-molecule and site-specific levels, facilitating the robust synthesis of per-read probabilities into site-wise modification scores.

Here we present WattmaMod, a label-efficient and lightweight framework for single-molecule nanopore DRS analysis that integrates wavelet-based multi-scale decomposition with contrastive pretraining and task-specific fine-tuning. By combining self-supervised representation learning on large-scale unlabeled signals with low-label supervised adaptation, WattmaMod is designed to capture transferable modification-associated patterns under limited annotation. To test its robustness and scalability for multi-modification detection, we systematically evaluated WattmaMod across RNA002 and RNA004 chemistries and diverse modification types. Specifically, under minimal labeled data conditions, WattmaMod achieves strong discriminative ability with an average AUC of 0.95 across seven RNA modification types, outperforming existing methods by an average of 10% and by up to 30% for specific types, demonstrating robust ranking performance even in low-resource settings. Further analyses in human cell lines, perturbation systems and cross-species settings suggested model-inferred potential co-occurrence patterns among RNA modifications, providing hypotheses for future experimental validation of coordinated RNA modification organization. These results establish WattmaMod as a scalable framework for accurate and extensible RNA modification profiling from nanopore DRS data.

## Results

### Overview of WattmaMod

To achieve accurate RNA modification detection from nanopore direct RNA sequencing (DRS) for both RNA002 and RNA004 chemistries, we developed WattmaMod, a unified learning framework that jointly models ionic current-level and event-level evidence. WattmaMod integrates raw current traces with event-level statistics—including mean, standard deviation, median, and dwell time—together with base-call quality scores and local sequence context to form a complementary representation for each candidate nucleotide site (Fig. 1a). This design directly targets two practical bottlenecks in DRS-based RNA modification detection: the often subtle modification-induced current perturbations and the long-range dependencies that are difficult to capture with single-scale signal representations. By coupling multi-scale signal encoding with sequence-context modeling, WattmaMod increases the separability of modified versus unmodified signals and improves robustness across datasets and chemistries (Fig. 1b,d). To further address limited supervision in many settings, we adopt a contrastive learning–based training strategy that combines self-supervised pretraining with supervised fine-tuning. This scheme yields more stable optimization and stronger generalization in few-shot regimes, with additional gains when paired with data augmentation (Fig. 1c,d; see Methods).

**Fig. 1.**
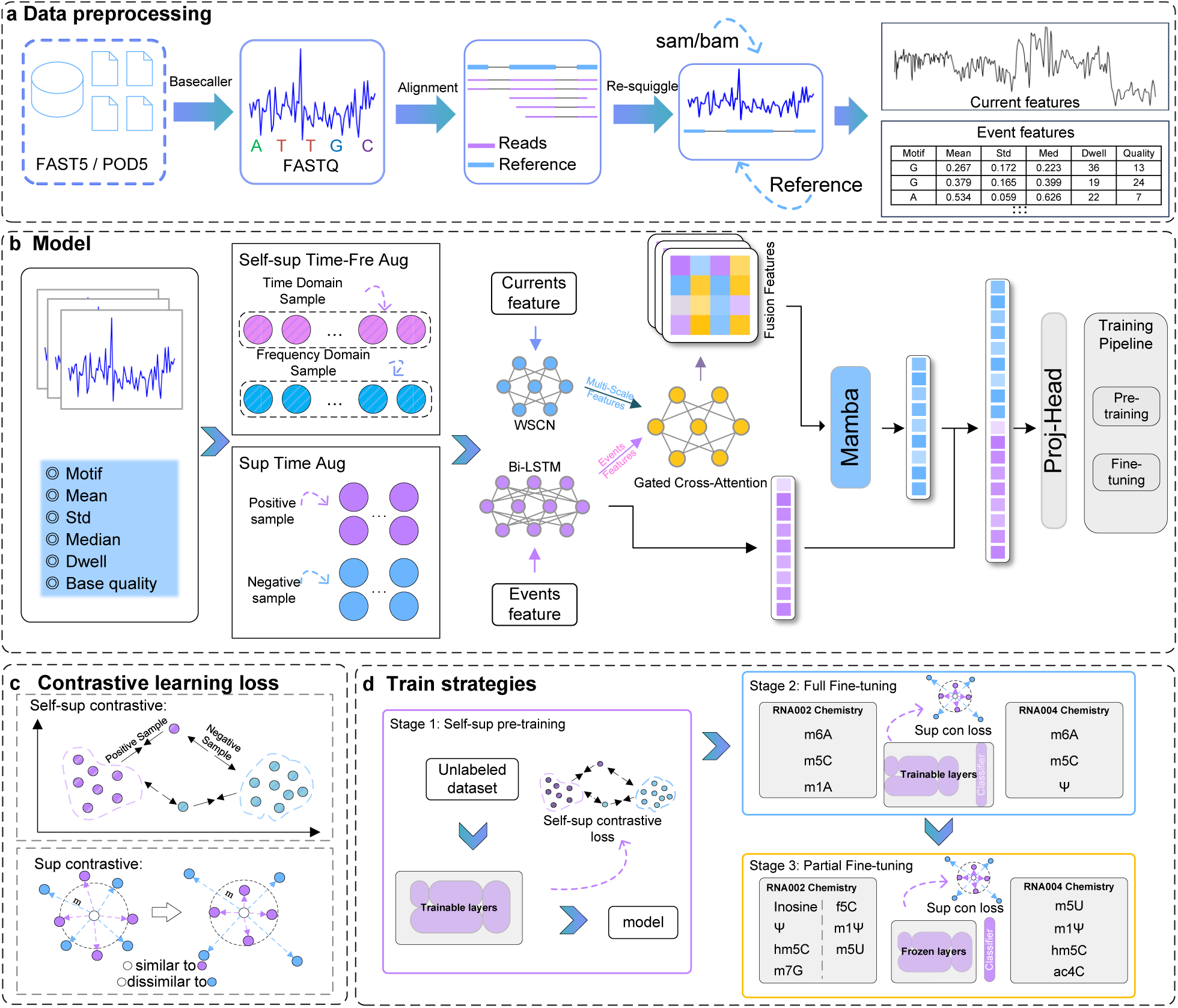
Overview of WattmaMod and data processing workflow. **a** Data preprocessing pipeline. Raw FAST5 and POD5 files are first basecalled to generate FASTQ sequences and aligned to the reference genome. Resquiggle/alignment is applied to obtain calibrated current signals and event-level features, including base identity, mean, standard deviation, median, dwell time, and base-call quality. **b** Schematic of the WattmaMod model architecture. The inputs, including normalized current signals and event descriptors, are processed by the WSCN and BiLSTM modules, fused via a gated cross-attention mechanism, and passed to a Mamba-based encoder for deep representation modeling. A projection head produces final embeddings for both pretraining and fine-tuning. **c** Contrastive learning losses. Self-supervised contrastive learning constructs positive and negative pairs through augmentation, while supervised contrastive learning leverages label information to promote intra-class compactness and inter-class separation. **d** Three-stage training strategy. Stage 1: Self-supervised pre-training on unlabeled data. Stage 2: joint full fine-tuning on abundant modifications from both RNA002 and RNA004 chemistries (RNA002: m6A, m5C, m1A; RNA004: m6A, m5C, Ψ). Stage 3: partial fine-tuning with encoder layers frozen for rare modification types (RNA002: inosine, Ψ, hm5C, m7G, f5C, m1Ψ, m5U; RNA004: m5U, m1Y, hm5C, ac4C).

To determine the optimal training configuration for RNA002, we benchmarked high-confidence in vitro resources, including the Curlcake synthetic sequence dataset ^38,39^, the ELIGOS in vitro transcription (IVT) dataset ^40^, and the in vitro epitranscriptomics (IVET) dataset derived from plant cDNA libraries ^29^. IVET with 5-mers achieved the best overall performance (Supplementary Fig. 4a,d,e), while training-resource choice strongly affected transferability: models trained on IVT reached ROC–AUC = 0.987 on matched data but dropped to 0.829 on IVET (Supplementary Fig. 4b,c), indicating that biological representativeness critically influences cross-dataset generalization. We therefore selected IVET with 5-mers as the default RNA002 configuration. For RNA004, the same 5-mer configuration was adopted considering the model architecture and training strategy, and Curlcake in vitro transcripts were further incorporated to provide higher-confidence supervision and test extensibility to the RNA004 chemistry ^41^. These results establish the training configuration used for subsequent WattmaMod evaluation and support its capacity to learn transferable modification-associated signals.

### Interpretable signal–feature remodeling driven by RNA modifications

To define the signal signatures captured by WattmaMod for modification discrimination, we systematically compared paired modified and unmodified reads from the IVT benchmark by jointly analyzing raw ionic current profiles, event-level statistics and feature attribution patterns. Across motifs, including the m6A-associated ACACU and the m5C-associated AACAG, modified reads were consistently separated from their unmodified counterparts in ionic current space, showing clear shifts in mean current and altered fluctuation patterns (Fig. 2a,b; Supplementary Fig. 6g). This perturbation extended beyond raw signals, modified reads exhibited reduced base-calling quality across multiple modification types, including m6A, m1A, m5C, hm5C, Ψ, m7G and inosine (Fig. 2c). At the event level, modification states were further associated with substantial reorganization of feature relationships, including distinct mean-variance, dwell-time and intensity coupling patterns under m5C and m6A, as well as modification-specific changes in variance and dwell time in A-centered contexts (Fig. 2d,f; Supplementary Fig. 2a). Feature attribution identified sequence context and base-calling quality as dominant predictors, with dwell time and signal mean providing additional support (Fig. 2e; Supplementary Fig. 2b). Because modified and unmodified IVT reads were generated from matched templates, the high attribution of sequence context is not primarily driven by sequence-composition imbalance, but instead reflects k-mer-dependent current perturbations, together with the contribution of base-calling quality to model weighting. Similar separation between modified and unmodified reads was maintained across datasets and modification types at the representation level (Supplementary Fig. 3), indicating that WattmaMod learns stable and coordinated signatures of RNA modification.

**Fig. 2.**
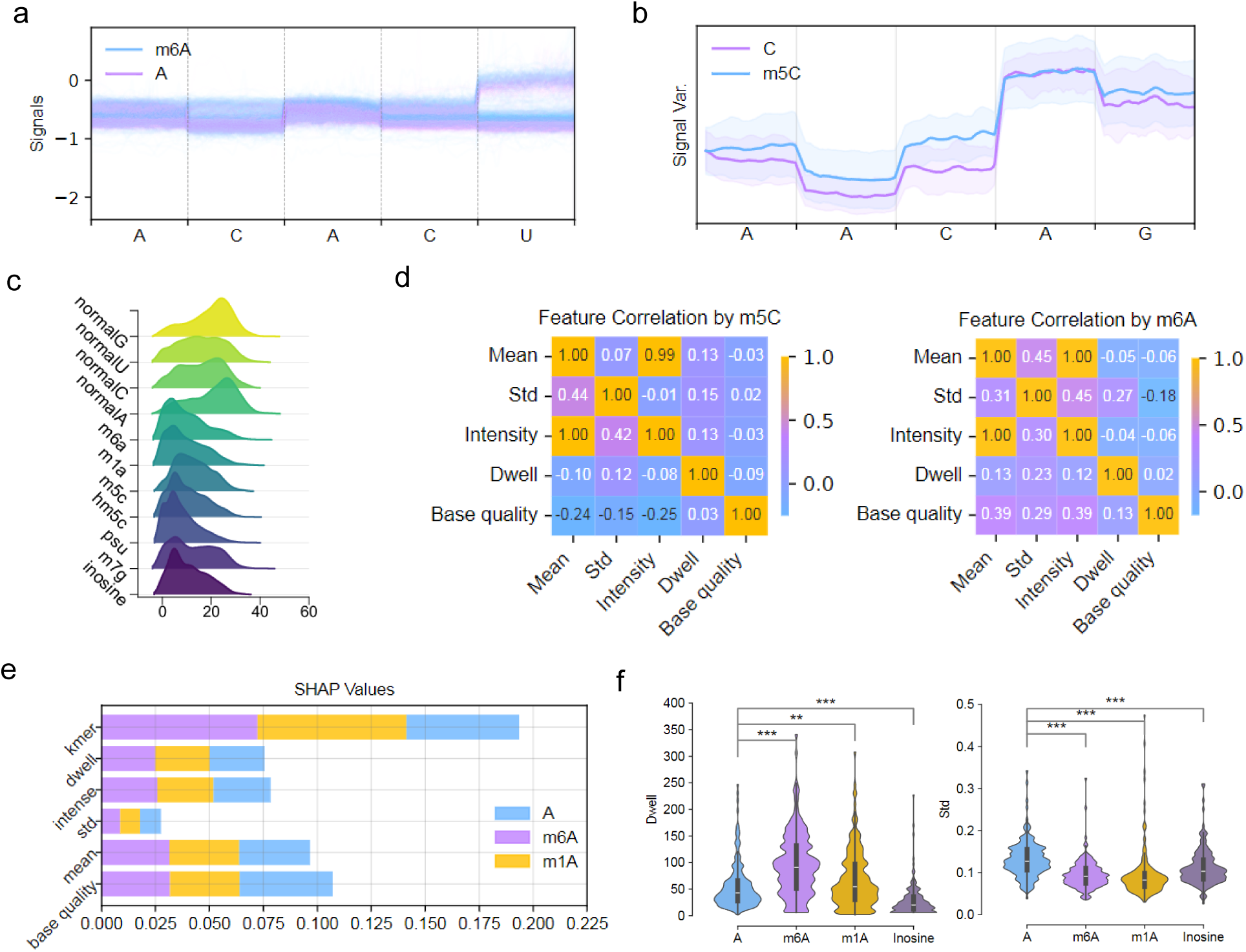
Distinct perturbation patterns induced by RNA modifications in DRS signals. **a** Comparison of normalized current signals for the 5-mer sequence *ACACU* between m6A-modified and unmodified adenosines. **b** Smoothed current profiles for the 5-mer sequence *AACAG* using a sliding window of 10 samples, showing amplitude deviations associated with m5C modification. **c** Ridge plots showing base quality distributions for modified nucleotides (m6A, m1A, m5C, hm5C, Ψ, m7G, inosine) and their unmodified counterparts (A, C, G, U). The x-axis denotes quality scores and the y-axis represents density. **d** Pearson correlation heatmaps of five event-level features, including mean, standard deviation, intensity, dwell time and base-calling quality, for m5C-modified samples (left) and m6A-modified samples (right). Correlations were calculated independently for each modification type, and color intensity indicates correlation strength. **e** SHAP summary bar plot showing the relative importance of five extracted features for distinguishing A, m6A and m1A. **f** Combined violin and box plots comparing dwell time and standard deviation for three adenosine modifications (m6A, m1A, A-to-I) versus unmodified A in the ELIGOS dataset. Statistical significance was assessed using two-sided Welch’s t-test (\**p <* 0.05, \*\**p <* 0.01, \*\*\**p <* 0.001).

### WattmaMod generalizes robustly across independent transcriptomes

WattmaMod generalized robustly across independent datasets and was further strengthened by interval-based dual-threshold confidence filtering under heterogeneous signal distributions ^30,42^. To test this, we evaluated the model on a non-overlapping IVET transcriptome. On the external IVET set, WattmaMod achieved ROC–AUC/PR–AUC values of 0.956/0.953 for m6A, 0.972/0.971 for m5C and 0.885/0.894 for m1A (Fig. 3a-c), improving m6A and m5C detection by approximately 2 percentage points over models trained under the same setting. F1 scores also remained consistently high across datasets under the default threshold (Supplementary Fig. 5a-e; Supplementary Fig. 6h). When dual-threshold confidence filtering was applied, performance further increased to 0.981/0.985 for m6A, 0.992/0.993 for m5C and 0.955/0.965 for m1A, while retaining approximately 75%, 81% and 63% of reads, respectively, at upper/lower thresholds of 0.9/0.3 (Fig. 3f). This gain was consistent with the clear bimodal separation of predicted probabilities between modified and unmodified reads (Fig. 3e). Read-length stratification further showed that performance increased from ∼0.87-0.89 in the 0-200 nt group to ∼0.93-0.97 for reads longer than 600 nt (Fig. 3d; Supplementary Fig. 5f), indicating stable behavior across read-length regimes. Compared with classical statistical and conventional machine-learning baselines, including logistic regression, GaussianNB, decision trees, random forests, KNN, SVM, XGBoost and multilayer perceptron, WattmaMod achieved the highest accuracy for both m5C (0.965) and m6A (0.913) (Supplementary Fig. 6i-j).

**Fig. 3.**
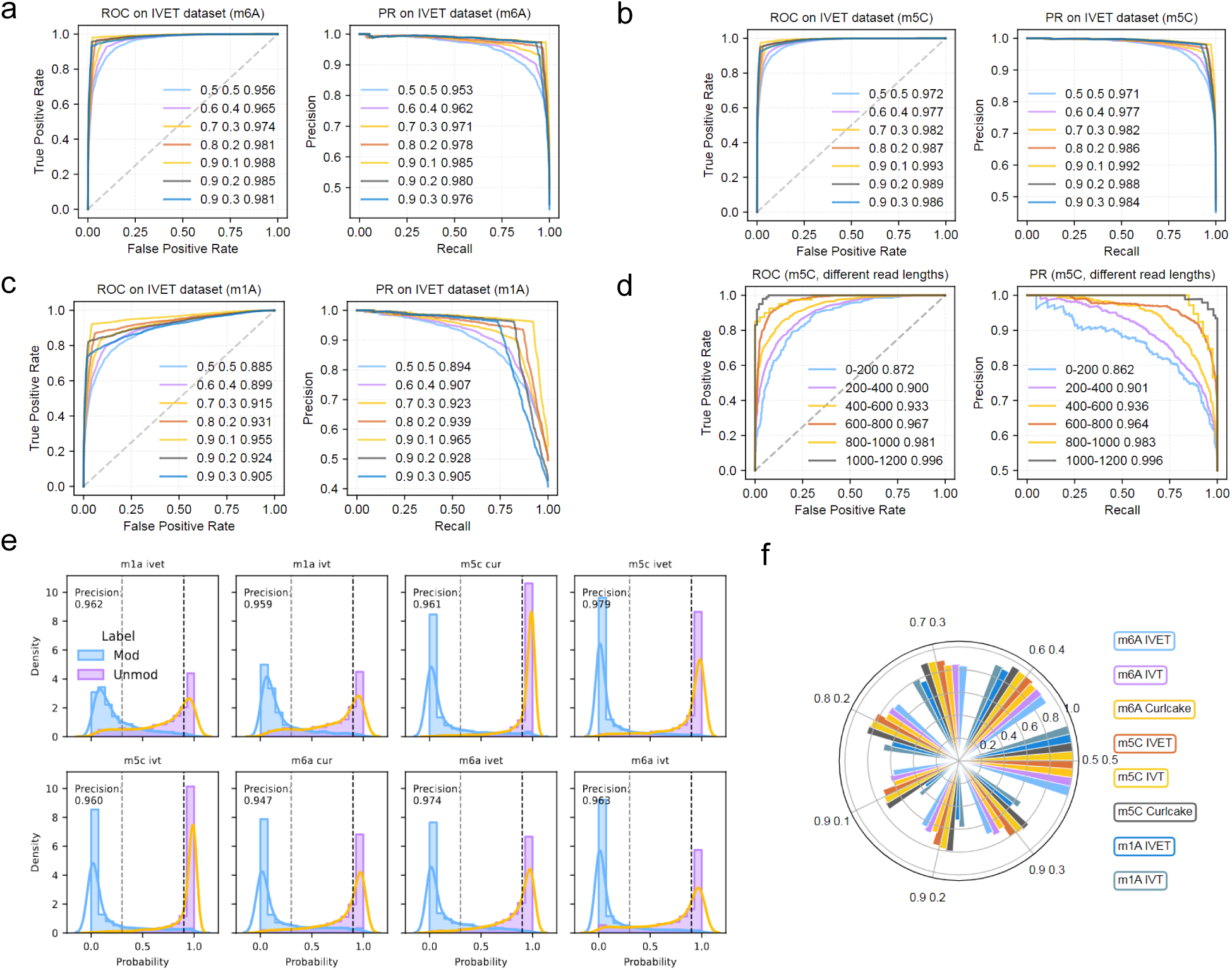
Cross dataset performance and confidence based improvement of WattmaMod. **a–c** ROC and PR curves for m6A (**a**), m5C (**b**), and m1A (**c**) classification on the IVET test set using models trained on IVET. Different colors indicate different interval based dual threshold settings (0.5/0.5, 0.6/0.4, 0.7/0.3, 0.8/0.2, 0.9/0.1, 0.9/0.2, 0.9/0.3). **d** ROC and PR performance of m6A detection across read length intervals (0–200, 200–400, 400–600, 600–800, 800–1000 and 1000–1200 nt) in the IVET dataset. **e** Density distributions of predicted probabilities for modified and unmodified reads under dual threshold confidence filtering. Probability separation is shown for m6A, m1A and m5C across the IVT, IVET and Curlcake datasets. Dashed lines indicate confidence thresholds, and precision is shown in each panel. **f** Retention rates of m6A, m5C and m1A under different dual threshold settings. Colors indicate different modification type and dataset combinations, and each segment shows the proportion of reads retained at the corresponding threshold.

### WattmaMod accurately detects multiple RNA modifications and quantifies stoichiometry

WattmaMod showed a clear overall advantage over existing tools across detection and stoichiometry benchmarks. To test this, we compared its performance in three settings, multi-modification benchmarking on IVT data, native-sample evaluation in K562 cells, and stoichiometry estimation using controlled mixing series. On the IVT benchmark, WattmaMod achieved the strongest overall performance across m6A, Ψ, m5C, inosine, m7G and m1A, with gains of ∼1 percentage point for m6A and up to ∼30 percentage points for hm5C relative to competing methods, although performance on m1A was slightly lower in selected settings (Fig. 4a). In transcriptome-wide native K562 nanopore DRS evaluation, where experimentally supported m6A-positive sites from MeRIP-seq ^43^ were greatly outnumbered by the vast background of non-modified transcriptomic sites, WattmaMod achieved a PR-AUC of 0.402, surpassing competing methods by 0.01 (Fig. 4b), indicating superior precision under biologically realistic class-imbalanced conditions. For quantitative stoichiometry estimation, WattmaMod achieved Pearson correlations of *r* = 0.952 for m5C and *r* = 0.932 for m6A (Fig. 4c-d), surpassing the best competing methods by ∼0.06 and ∼0.03, respectively.

**Fig. 4.**
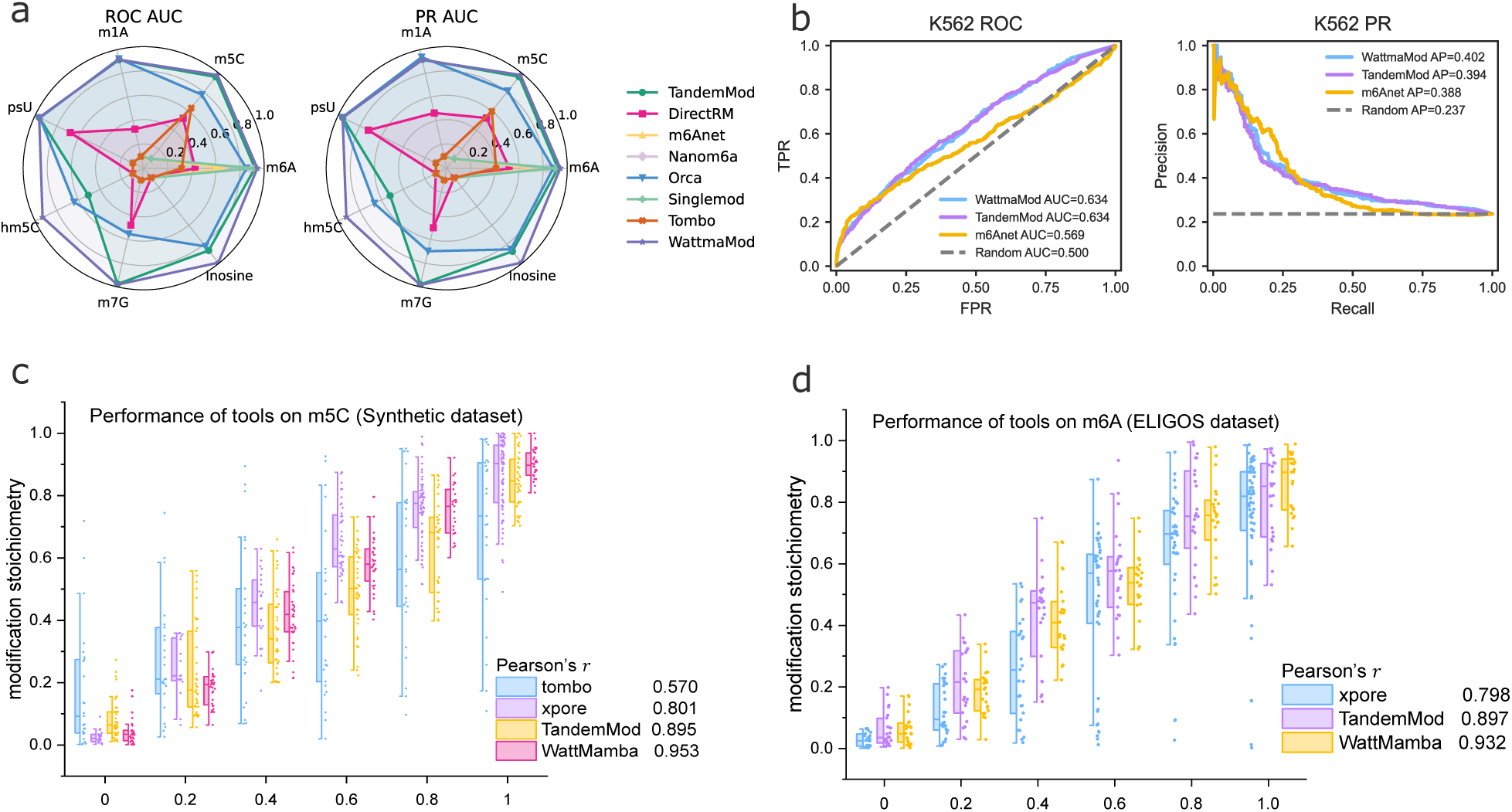
Comparison of WattmaMod and existing tools for RNA modification detection and stoichiometry estimation. **a** Radar plots summarizing detection performance across multiple tools on the ELIGOS in vitro benchmark for different RNA modification types, including m6A, m5C, m1A, m7G, hm5C, Ψ and Inosine. ROC-AUC values are shown on the left and PR-AUC values on the right. **b** Site level evaluation in K562 cells using MeRIP-seq as orthogonal reference. ROC curves (left) and PR curves (right) are shown for m6A prediction by WattmaMod and representative methods. The diagonal dashed line indicates random performance. **c** Quantitative stoichiometry estimation on a synthetic m5C dataset. Predicted stoichiometries are shown across ground truth stoichiometry bins for Tombo, xPore, TandemMod and WattmaMod. Box plots show the interquartile range with the median as the center line, and points indicate individual predictions. Pearson’s correlation coefficien (*r*) is shown to assess agreement with the ground truth. **d** Quantitative stoichiometry estimation on the ELIGOS m6A dataset for xPore, TandemMod, and WattmaMod. Pearson’s correlation coefficient (*r*) indicates the consistency between predicted and true stoichiometries.

### Multi-stage training enables low-label expansion across multiple RNA modifications and sequencing chemistries

We next investigated whether the multi-stage training strategy enables WattmaMod to achieve accurate low-label expansion across diverse RNA modifications and nanopore sequencing chemistries, including RNA002 and RNA004. To do this, we evaluated the model on RNA002 and RNA004 using only 3,000 positive and 3,000 negative samples per modification for fine-tuning. Under RNA002, WattmaMod achieved strong performance across multiple modification types, with PR-AUC values of 0.968 for f5C, 0.929 for hm5C, 0.998 for inosine, 0.946 for m1A, 0.921 for m5U, 0.968 for m7G and 0.989 for Ψ, while also maintaining similarly strong performance for m5U and m1Ψ in Curlcake (Fig. 5a). Lift analysis further showed that the top-ranked 10-30% of read segments recovered most true sites (Supplementary Fig. 8), indicating effective prioritization even with minimal labeled data. We further evaluated whether this multi-modification capability could be retained under RNA004. In this setting, modified and unmodified reads remained clearly separable, F1 scores exceeded 0.94 across most coverage strata (Fig. 5b; Supplementary Fig. 9), and false positive rates remained low across modification types in both RNA002 (0.042–0.127) and RNA004 (0.038–0.114) (Supplementary Fig. 20a). Dorado RNA004 modified-base calling was further included as a reference on the Curlcake synthetic data, using matched modified and unmodified samples to generate PR curves for m6A, m5C and Ψ (Supplementary Fig. 20c). After fine-tuning on the RNA-modbase IVT dataset to adapt to the shorter effective signal context of RNA004, WattmaMod achieved ROC-AUC values of 0.983-0.999 and PR-AUC values of 0.976-0.999 for m6A, inosine, Ψ and m5C (Supplementary Fig. 12b,d), indicating that the model is readily transferable beyond a single sequencing chemistry. The benefit of multi-stage training was further supported by training analyses. Fine-tuning with 3,000 samples yielded the best overall performance (Supplementary Fig. 19), and pretraining modestly improved all validation metrics (Supplementary Fig. 20b). t-SNE showed clearer separation between modified and unmodified reads after fine-tuning (Fig. 5c; Supplementary Fig. 6a-f), and pretraining provided an additional ∼1-2 percentage-point gain in validation accuracy together with faster convergence (Fig. 5d). Overall, multi-stage training improved low-label adaptation efficiency.

**Fig. 5.**
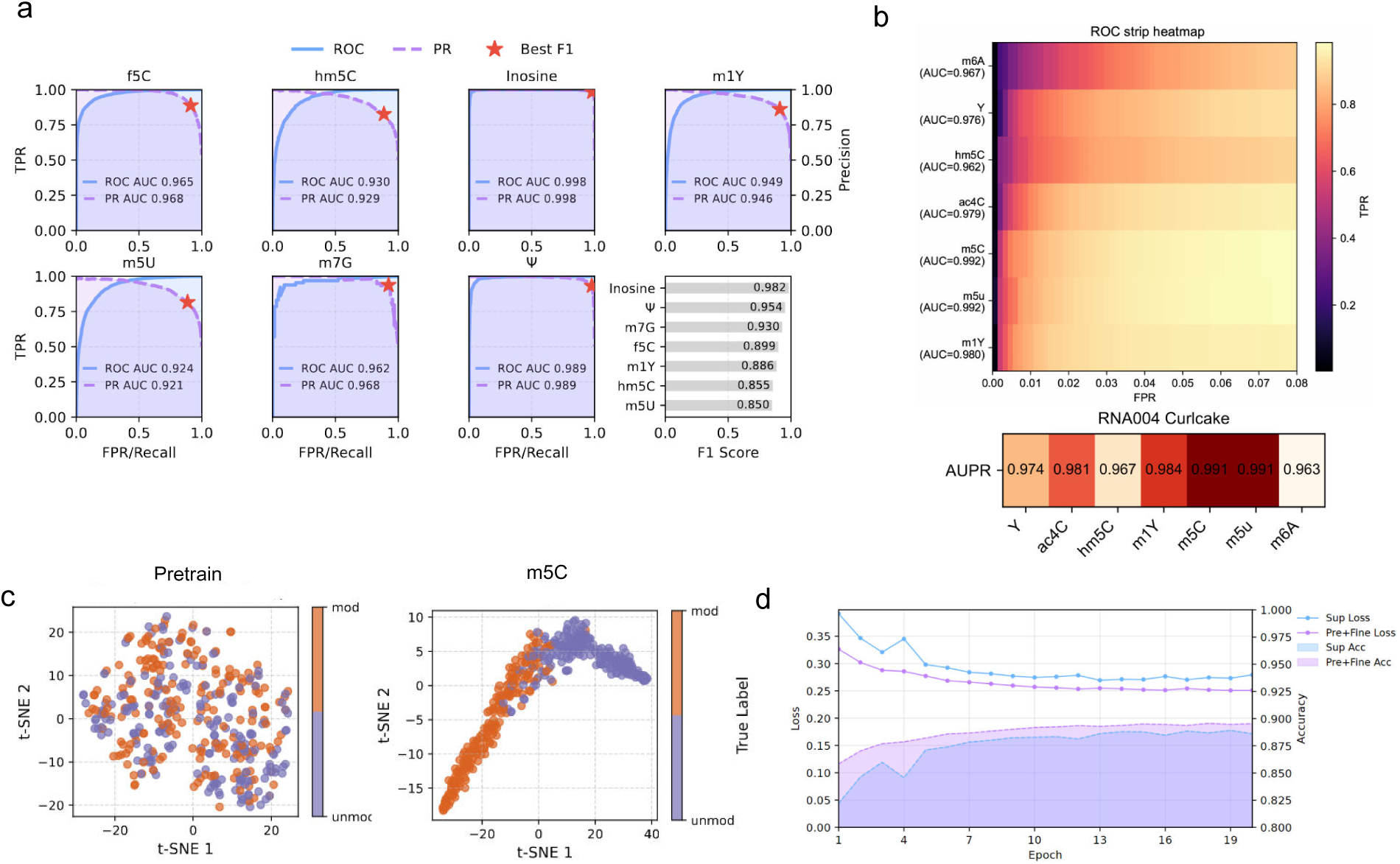
Multi stage training enables low label expansion across multiple RNA modifications and nanopore chemistries. **a** ROC and PR curves showing detection performance across multiple RNA modification types, including hm5C, inosine, Ψ, f5C, m5U, m1A, m6A, m1Ψ and m7G. ROC-AUC, PR-AUC and optimal F1 scores are shown for each modification. **b** Performance of RNA modification detection on the RNA004 Curlcake in vitro transcription dataset, shown as a ROC strip heatmap (top) and an AUPR heatmap (bottom). **c** t-SNE visualization of embedding representations learned during self supervised pretraining (left) and after supervised fine tuning (right), showing improved separation between modified and unmodified reads after fine tuning. **d** Line plots show the training loss for the pretrained and fine tuned model (orange) and the directly supervised model without pretraining (blue). Shaded regions indicate validation accuracy during training

### Conserved and cell line specific epitranscriptomic landscapes across human cell lines

We next assessed whether WattmaMod can accurately characterize RNA modification landscapes across heterogeneous human cell lines. To do this, we analyzed six tumor cell lines (A549, HCT116, MCF7, HEPG2, K562 and HEYA8) together with H9 embryonic stem cells and HEK293T. High-confidence m6A sites predicted by WattmaMod were strongly enriched in the canonical DRACH motif, with GGACT and AGACT as the dominant 5-mer contexts (Fig. 6a; Supplementary Fig. 20a). WattmaMod also identified non-DRACH high-confidence sites, including GGACG and AGACG, whose relative abundance varied across cell types. Predicted m6A sites were further enriched near the CDS–3^′^UTR junction and across the 3^′^UTR, with a broad signal extending ∼1,000 nt upstream of transcript ends (Supplementary Fig. 10a,b). These patterns indicate that WattmaMod recapitulates known m6A sequence and positional features in complex transcriptomes. Comparison with HepG2 m6A-seq data showed that 70% of the 5,680 genes predicted by WattmaMod were supported by independent experimental evidence (Fig. 6c). In HEYA8 cells, most modified transcripts contained a single high-confidence modification site (Fig. 6b). Among randomly selected genes, ACTB, PTMA, RPLP1 and RPS13 showed relatively high m6A levels, whereas FTL, Hsp70 and RPS26 displayed cell-line-dependent patterns (Fig. 6e), consistent with known m6A transcriptome landscapes ^44–46^. In H9 embryonic stem cells, gene-level profiling revealed highly regulated loci consistent with canonical m6A regulation (Supplementary Fig. 22). In HEK293T, m6A-modified genes were significantly enriched in RNA processing, splicing, RNA transport and ubiquitin-related pathways (Supplementary Fig. 10d; Supplementary Fig. 11). WattmaMod also enabled characterization of multiple RNA modification types. Motif maps recovered expected sequence contexts for m6A in HEYA8 and for m5C in WT/NSUN2-KO HeLa cells, while motif-logo analysis across ten RNA modifications revealed distinct sequence preferences in mRNA and non-coding RNA contexts (Supplementary Fig. 21b-d).

**Fig. 6.**
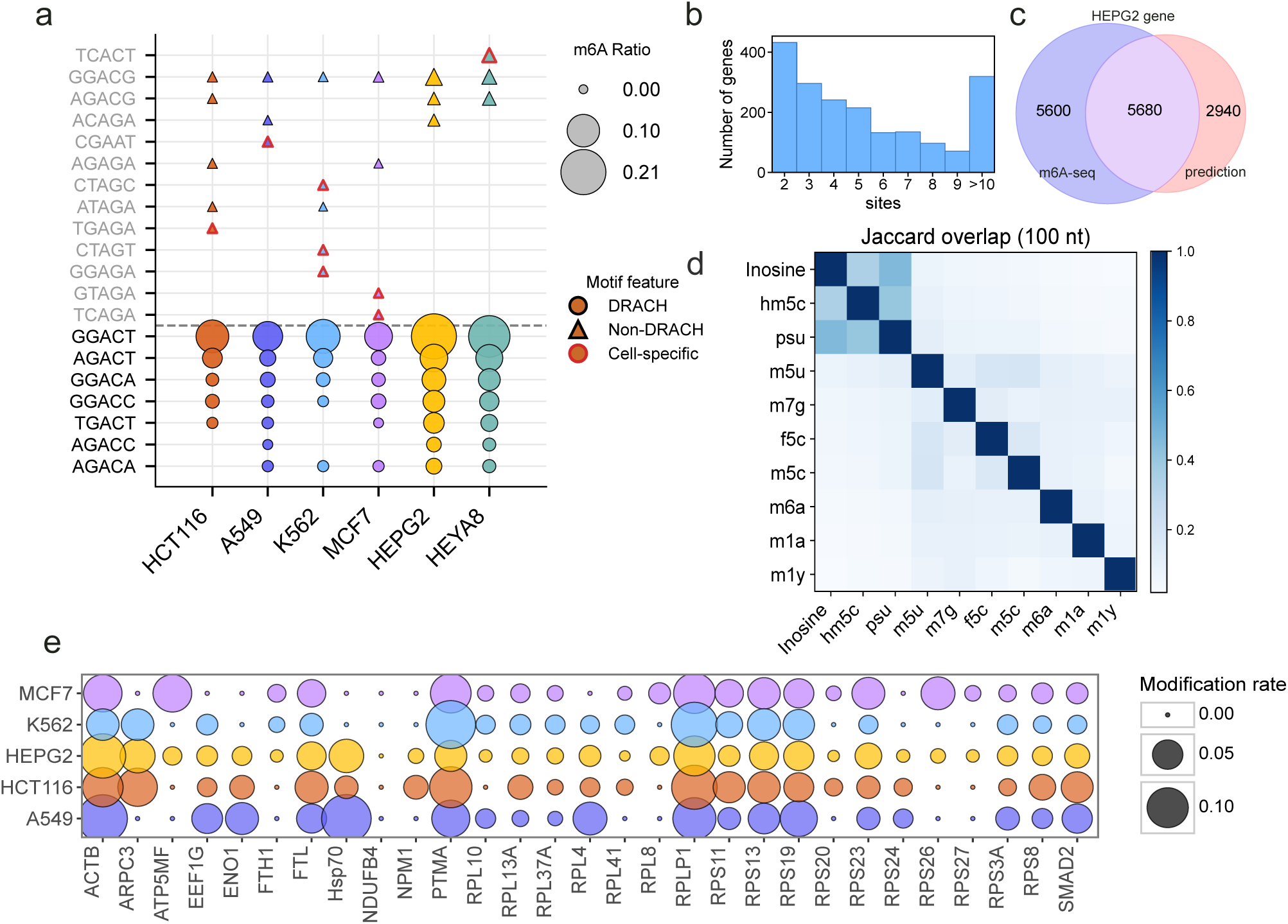
Transcriptome wide RNA modification landscapes across human cancer cell line. **a** Top 10 enriched m6A associated 5-mer motifs across six human cancer cell lines, including MCF7, K562, HCT116, HEPG2, A549 and HEYA8. **b** Distribution of the number of modified sites per gene in HEYA8 cells. **c** Overlap between the m6A modified gene set predicted by WattmaMod and the gene set identified by m6A-seq. Blue indicates genes detected by m6A-seq, red indicates genes predicted by WattmaMod, and the overlapping region indicates genes identified by both methods. **d** Predicted local co-modification landscape of ten RNA modification types in HEYA8 cells, quantified by pairwise Jaccard overlap within a ±100-nt window. The heatmap shows pairwise overlap between inosine, hm5C, Ψ, m5U, m7G, f5C, m5C, m6A, m1A and m1Ψ site sets. Two sites were considered co localized when their transcript coordinates were within ±100 nt. Darker colors indicate higher overlap, and the diagonal indicates self overlap. **e** Bubble plot showing the distribution of m6A modification levels in 30 randomly selected genes across five human cancer cell lines. Circle size indicates modification intensity, and color denotes cell line identity.

### Cross-species perturbation-responsive RNA modification dynamics and predicted higher-order multi-modification organization

A key strength of WattmaMod is that it sensitively captures biologically expected RNA modification changes across species and perturbation settings, extending its applicability beyond static single-condition detection. To evaluate this, we tested WattmaMod in genetic perturbation datasets from human, Arabidopsis, and yeast. Across all settings, WattmaMod consistently recovered the expected direction of change, including a marked reduction of m6A in METTL3-KO HEK293T cells (median: 0.11 to 0.06; Fig. 7a) and a decrease of m5C in NSUN2-KO HeLa cells (median: 0.13 to 0.08; Supplementary Fig. 13). Similar perturbation-associated shifts were also detected for m6A in Arabidopsis mutants/vir-c and for modification variation in yeast rRNA (Fig. 7b; Supplementary Fig. 14;Supplementary Fig. 17a-d). In addition, WattmaMod showed high technical reproducibility, with cross-replicate Pearson correlations of 0.996 in H9 and 0.954 in HEYA8 (Supplementary Fig. 18).

**Fig. 7.**
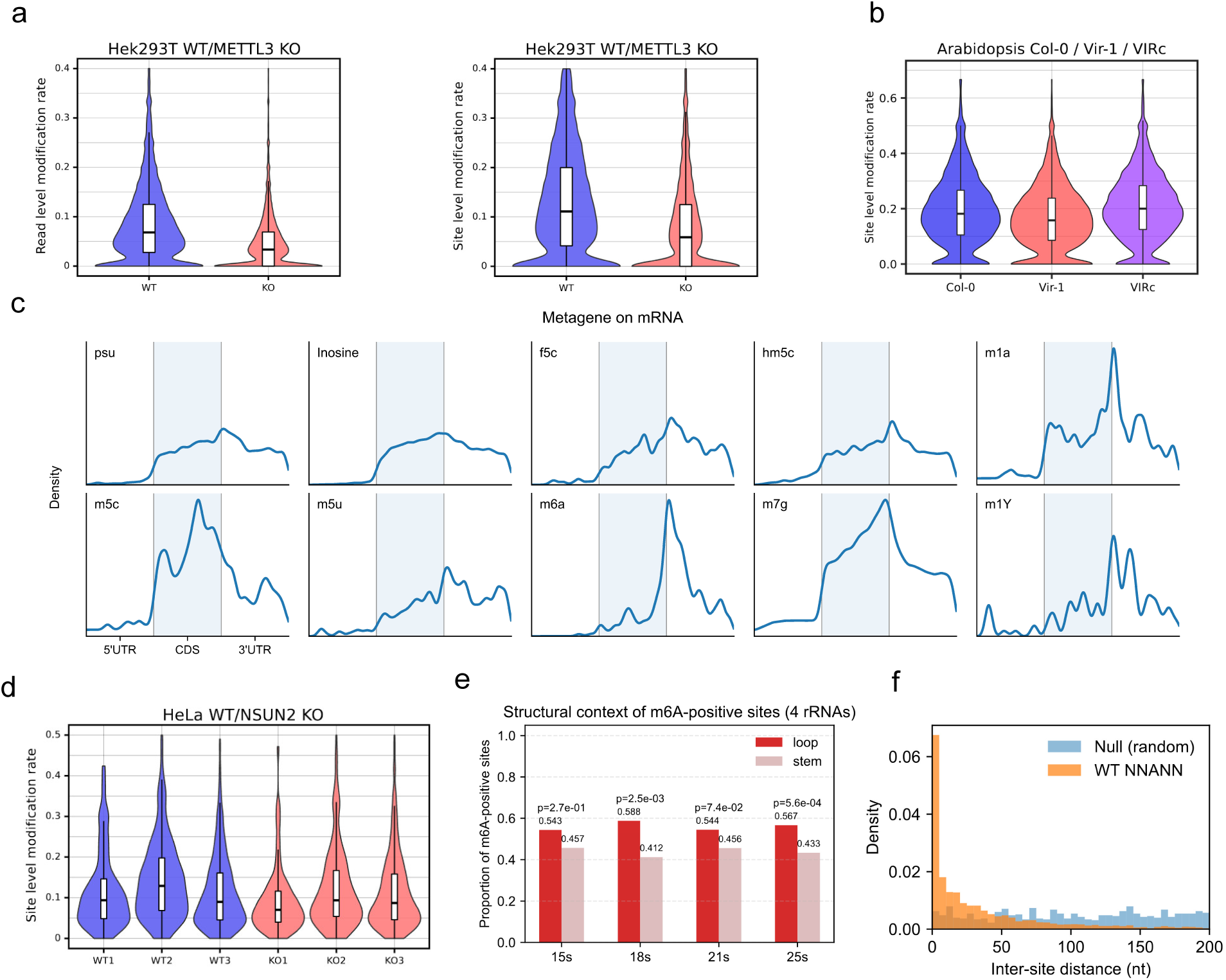
Cross-species validation of perturbation-responsive RNA modification patterns and spatial organization. **a** Violin plots of m6A modification probabilities in wild type and METTL3 knockout HEK293T cells. Embedded box plots indicate the median, interquartile range and whiskers extending to (1.5×IQR). **b** Site level distributions of m6A modification rates in Arabidopsis wild type Col-0, the VIR mutant vir-1 and the complemented line vir-c. **c** Metagene density profiles of ten RNA modifications across normalized transcript regions (5^′^UTR, CDS and 3^′^UTR). Shading indicates the CDS, and vertical lines mark region boundaries. **d** Violin plots comparing site level m5C modification rates in wild type HeLa cells and NSUN2 knockout cells. **e** Structural classification of high confidence m6A sites on *Saccharomyces cerevisiae* 15S, 18S, 21S and 25S rRNAs as loop or stem regions based on RNAfold predicted secondary structures. Bar plots show loop and stem proportions, and P values were calculated using two tailed binomial tests under the null hypothesis of equal loop and stem proportions. **f** Nearest neighbor distance distributions of high confidence m6A sites on *Saccharomyces cerevisiae* rRNAs (orange) compared with randomly sampled control sites from the same transcripts (blue).

A notable application of WattmaMod is its ability to generate model-based inferences of potential local co-occurrence patterns among distinct RNA modifications, moving beyond conventional site-by-site prediction. Meta-gene analysis across ten RNA modifications supported transcript-level spatial characterization within a unified analytical framework (Fig. 7c). Among them, the m1A distribution should be interpreted with caution because m1A prediction showed lower reliability than other modifications and may be partially affected by m6A-related signal interference. Notably, the model predicted a potential local co-occurrence pattern involving Inosine, hm5C, and Ψ in HEYA8, with multiple pairwise Jaccard indices greater than 0.2 within a ±100-nt window (Fig. 6d). A similar but weaker pattern was also observed in wild-type HEK293T, where local overlap values were around 0.1 (Supplementary Fig. 15), suggesting that this predicted three-modification association may recur across cellular contexts. In *Populus tridentata*, we further detected substantial m6A/m1A site overlap (Jaccard = 0.375) despite negligible rate-level correlation (*R*^2^ = 0.01), indicating that strong local spatial association can exist even in the absence of global quantitative concordance (Supplementary Fig. 16). Based on these observations, we refer to such localized regions of recurrent multi-modification overlap as putative multi-modification hubs.

Orthogonal structural analysis further supported the biological plausibility of the modification landscapes identified by WattmaMod. To examine whether predicted sites were consistent with known structural preferences, we aligned site-level predictions with RNA secondary structure annotation in yeast rRNA. RNAfold-based analysis ^47^ showed that high-confidence m6A sites were significantly enriched in loop regions and displayed stronger local clustering than randomized controls (*P* = 0.027 to 5.6 × 10^−5^; Fig. 7f,g; Supplementary Fig. 17e). These results provide independent support that WattmaMod-derived modification patterns are consistent with established structural constraints.

## Discussion

To address major challenges in DRS-based modification detection—including limited high-confidence labels, heterogeneous data quality, the difficulty of jointly profiling multiple modification types, and limited interpretability—we present WattmaMod, a deep-learning framework for RNA modification detection. With supervised fine-tuning on only 250,000 labeled single-molecule reads, WattmaMod outperforms widely used methods in both detection accuracy and inference throughput, while substantially accelerating GPU execution and reducing computational cost. Owing to its label-efficient and few-shot adaptability, WattmaMod maintains stable cross-dataset generalization and supports diverse modification types.

WattmaMod adopts multiscale signal encoding guided by discrete wavelet transform (DWT), where low-frequency components capture background and steady-state structures and high-frequency components emphasize transient perturbations introduced by modified bases, enabling selective extraction of modification-related cues across frequency bands. Dynamic cross-attention further leverages event-derived priors to adaptively reweight multi-scale channels. In addition, the state-space module integrates long-range context, allowing signal and sequence features to form a context-aware joint representation. Training follows a two-stage contrastive paradigm: self-supervised pretraining constructs time–frequency–enhanced positive/negative pairs to obtain noise-robust embeddings, followed by supervised contrastive fine-tuning with limited labels, where amplitude and temporal perturbations improve class separation and generalization. Together, band decomposition, prior-guided adaptive modulation, SSM-enabled long-range dependency modeling, and self-supervised initialization contribute to improved robustness, generalization and overall detection performance relative to baseline approaches.

Across multiple in vitro benchmarks, WattmaMod shows improved detection performance across diverse modification types compared with existing tools. It further demonstrates stable cross-species generalization in human tumour cell lines, poplar, *Arabidopsis* and *Saccharomyces cerevisiae*. The model enables high-precision single-molecule detection and reliably captures changes in modification levels across genetic perturbation backgrounds, indicating that WattmaMod learns transferable biophysical signal patterns rather than dataset-specific noise.

Despite these strengths, several limitations remain. First, some training and evaluation resources—such as Curl-cake long transcripts generated under RNA004 chemistry and RNA-modbase synthetic oligonucleotides with BED-defined modification sites—deviate from native transcriptomes in biological context, structural complexity and read-length distributions. Combined with reduced signal-to-noise near fragment ends and less stable event alignment, these factors partially impair performance on short reads. In practice, longer reads should be prioritized during inference, together with stringent coverage filtering to obtain higher-confidence sites. Second, model performance depends strongly on training representativeness. IVET-trained models show markedly improved cross-species and cross-platform generalization compared with IVT-trained counterparts, suggesting that diversity of modification contexts and sequence environments is often more important than simply scaling dataset size. Third, the current model does not explicitly incorporate secondary-structure priors during inference, which is known to modulate modification accessibility and signal manifestation, particularly for structure-sensitive marks. Incorporating structure-aware features (for example, pairing probability, accessibility, or loop/stem annotations derived from RNAfold-like predictors) may further improve site resolution and robustness in complex transcriptomic contexts. Future work will incorporate broader species and tissue sources to build a more neutral and representative foundation dataset, while integrating secondary-structure priors to mitigate synthetic bias and enhance biological fidelity when interpreting real samples.

WattmaMod also provides a scalable basis for a unified multi-task framework. Such a framework could enable parallel detection of dozens of modifications within a single model, while integrating spatiotemporal visualization and rule extraction to characterize competitive and cooperative relationships and further improve interpretability. From an application perspective, WattmaMod allows rapid adaptation to specific sample types or target marks using only a small set of validated sites, enabling efficient high-throughput inference for identifying disease-associated or condition-specific RNA modification programs.

## Methods

### Data and preprocessing

For direct RNA sequencing (DRS) data generated with the RNA002 kit, reads were basecalled using Guppy v6.1.5 with the RNA high-accuracy model. Multi-read FAST5 files were then converted to single-read FAST5 format using ont fast5 api v4.0.0. The basecalled reads were aligned to the reference transcriptome using minimap2 v2.22^48^. Raw current signals were subsequently processed with the *resquiggle* procedure in Tombo v1.5.1^49^ to obtain event-level segmentation and accurate signal-to-reference alignments.

For RNA004 chemistry, POD5 files were basecalled using Dorado v0.9.6^50^. Raw current signals and sequences were then jointly aligned to the reference transcriptome using Uncalled4 v4.1.0^51^, producing BAM files annotated with signal index tags (ul and ur). On this basis, we implemented custom scripts to extract raw signal values at single-nucleotide resolution by combining BAM alignments with signal index tags. When the alignment model produced invalid signal indices (e.g. ul = 0), we applied a forward-filling strategy that replaces them with the nearest preceding valid index, thereby preserving continuity of the signal–base mapping without introducing substantial bias. Insertions and deletions in the CIGAR string were handled by inserting gap characters (“-”) into the reference or read sequences, respectively, and padding the associated signal segments to maintain a one-to-one positional correspondence between aligned bases and signal samples. Command-line options for these steps are in Supplementary Table 1.

During signal feature construction, we normalized the raw current traces to reduce systematic bias, segmented signals using Tombo or Uncalled4, and spline-interpolated each segment to a fixed length of 500 samples. For each event, we computed the mean, standard deviation, median, dwell time, and base-call quality. We then used a 5-mer window centered on the target nucleotide to aggregate the event-level statistics with the normalized signal features, forming the input to the deep learning model(Fig. 1a).

### Training strategy for models

Direct end-to-end supervised training is prone to overfitting when confronted with multiple RNA modification types and sparse annotations, which in turn limits the quality of learned representations. We adopt a multi-stage training strategy. We first perform self-supervised pretraining on a blended RNA002 dataset using only unlabeled, read-level samples (2.5 million) from real HEK293T data. Through instance-wise contrastive learning, the model learns general biological representations without relying on annotation labels.

After obtaining the general encoder, the second stage conducts supervised fine-tuning on labeled read-level data using a distance-based supervised contrastive objective to establish stable discriminative boundaries. We performed full-parameter supervised fine-tuning across the primary modifications in both chemical systems (RNA002: m6A, m5C, m1A; RNA004: m6A, m5C, Ψ). This stage-wise grouping was defined mainly according to the availability, scale, and reliability of high-confidence labeled training data for each modification type under each nanopore chemistry, with biological abundance considered as an additional factor influencing data accessibility.

The third stage targets low-resource and hard-to-obtain modification types by unfreezing the terminal projection layer and classification head while keeping the backbone encoder parameters unchanged, enabling rapid incremental adaptation for each new classification task. RNA002 focuses on Inosine (A-to-I), Ψ, hm5C, m7G, f5C, m1Ψ, and m5U; RNA004 focuses on m5U, m1Ψ, hm5C, and ac4C.

### Self-supervised contrastive learning in the time–frequency domain

During unlabeled pretraining, we constructed dual-view contrastive samples in the time and frequency domains and optimized their consistency using the NT-Xent ^52^ objective. This strategy encouraged the model to learn representations robust to signal non-stationarity and spectral perturbation. To broaden the coverage of augmentation, we combined standard time-domain transformations with frequency-domain perturbation operators, thereby improving robustness to local distortion, spectral variation and background noise in nanopore signals.

For each input signal *x_i_*, one time-domain augmentation was randomly selected from dynamic jitter, amplitude scaling and local temporal reordering to generate the time-domain view **v***_T_*, whereas the frequency-domain view **v***_F_* for the same sample was defined as:

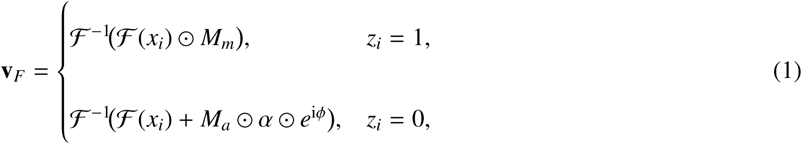

where F and F ^−1^ denote the Fourier transform and its inverse, respectively. Here, *z_i_* ∈ {0, 1} is a random indicator for spectral masking or spectral perturbation. Specifically, *M_m_* ∼ Bernoulli(1 − *p_m_*) denotes the frequency-retention mask, *M_a_* ∼ Bernoulli(*p_a_*) denotes the frequency-addition mask, *ϕ* ∼ U(0, 2*π*) controls the phase, and *α* ∼ U(0, 0.1 |F (*x_i_*)|) controls the perturbation magnitude.

To leverage unlabeled RNA sequencing data under limited annotation, the time-domain and frequency-domain views (**v***_T_,* **v***_F_*) generated from the same sample were treated as a positive pair, whereas views from other samples within the same mini-batch were treated as negative pairs, and the model was optimized using the NT-Xent loss with temperature *τ* = 0.15:

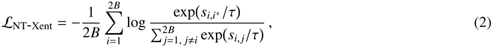

where *s_i j_* denotes the cosine similarity between samples *i* and *j*, *i*^+^ denotes the positive counterpart of sample *i*, and *B* is the batch size. This objective encouraged the model to learn stable representations from unlabeled signals and provided robust initialization for subsequent supervised fine-tuning (Supplementary Fig. 7a,d).

### Composite augmentation for supervised fine-tuning

During supervised fine-tuning, we applied composite augmentation to labeled samples to mimic amplitude fluctuation and nonlinear temporal distortion in nanopore signals, thereby improving robustness while generating multiple views for contrastive supervision. For each input signal *x*(*t*), we first placed *K*+2 control points 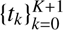 uniformly over the time domain, with the default setting *K*=2. Amplitude scaling factors 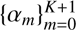 were then independently sampled at these control points from 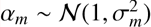, and cubic-spline interpolation converted them into a continuous modulation curve *g_c_*(*t*), whose pointwise multiplication with *x*(*t*) produced an amplitude-warped signal *y*(*t*). At the same control-point locations, temporal scaling factors 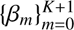 were independently sampled from 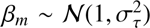, and cubic-spline interpolation yielded a continuous temporal scaling curve *β_c_*(*t*). To preserve the valid signal range and the original sequence length, we defined the normalized temporal mapping as:

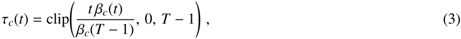

where *T* denotes the sequence length. The final augmented signal 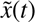 was obtained by linearly resampling *y*(*t*) along *τ_c_*(*t*) (Supplementary Fig. 7b,e). In this way, the augmentation sequence first applied amplitude warping and then temporal warping, which more closely mimicked gain drift and temporal stretching in practical nanopore signals.

The augmented samples were further used to construct supervised contrastive pairs during fine-tuning. Specifically, samples with the same label were treated as positive pairs, whereas samples with different labels were treated as negative pairs, and the embeddings were optimized using a pairwise distance-based contrastive loss:

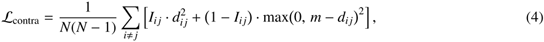

where *I_i j_* = 1 if samples *i* and *j* share the same label and *I_i j_* = 0 otherwise, *d_i j_*denotes the cosine distance between their embeddings, and *m >* 0 is the margin imposed on negative pairs. Positive pairs were therefore pulled together by the quadratic term, whereas negative pairs were pushed apart by the hinge term.

The final optimization objective combined the cross-entropy loss with the supervised contrastive term and was written as:

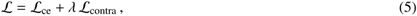

where *λ* was set to 0.8 and L_ce_ denotes the cross-entropy loss. This objective improved discrimination while maintaining robustness to amplitude and temporal perturbations as well as batch-level variation.

### Wavelet-guided multi-scale learnable encoder

To capture the non-stationary characteristics of nanopore sequencing signals across multiple temporal scales, we employed discrete wavelet decomposition using the Daubechies-4 wavelet with *J* = 4 levels, followed by learnable channel reweighting and one-dimensional convolution for feature integration. For an input signal *x* ∈ R*^B^*^×*T*^, the wavelet transform 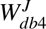 produced a set of approximation and detail coefficients:

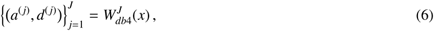

where *a*^(J)^ denotes the low-frequency approximation component and 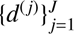 denote the detail components at different scales (Supplementary Fig. 7c).

To preserve the physical characteristics of different frequency bands while enabling unified downstream processing, each scale component was resampled to a fixed length *L*^∗^ = 200. Specifically, first-order linear interpolation was applied to the low-frequency component to obtain 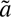, whereas nearest-neighbor interpolation was applied to each high-frequency component to obtain 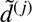, thereby preserving abrupt local changes without introducing additional smoothing. The resampled components were then concatenated along the channel dimension to form the multi-scale representation 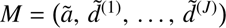.

To adaptively model the relative contributions of different frequency bands, we introduced a learnable channel reweighting module, which produced a reweighted representation *M*^′^ as:

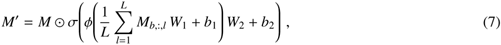

where *ϕ* and *σ* denote the ReLU and Sigmoid functions, respectively, *W*_1_ ∈ R*^C^*^×*d*^ and *W*_2_ ∈ R*^d^*^×*C*^ are learnable projection matrices, with *C* = 5 and *d* = 16, and the resulting channel-wise weights were broadcast along the temporal dimension. In this way, the model dynamically emphasized informative scales while suppressing less relevant components.

The reweighted tensor *M*^′^ was then fed into a convolutional encoder composed of two one-dimensional convolutional layers and one max-pooling layer with ReLU activations, yielding the signal feature representation *X*_sig_ ∈ R*^B^*^×*D*×*L*′^ for downstream modules. Boundary handling was implemented using the wavelet zero mode to avoid reflection artifacts, and both the multi-scale decomposition and channel reweighting remained fully differentiable during training.

### Dynamic cross-attention and sequence modeling

To address the conflicts and noise introduced by simple feature concatenation, we used an adaptive fusion strategy to integrate heterogeneous modalities and thereby combine local signal details with global event-level regularities. After BiLSTM encoding, the event-level statistical features were represented as *X*_stats_. We then used *X*_sig_ as the query and *X*_stats_ as the key and value, so that the signal representation could selectively retrieve complementary information from the statistical branch. The fused representation was defined as:

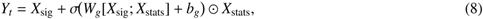

where *W_g_* and *b_g_* denote the learnable parameters of the gating network. This design enabled context-aware feature modulation by enhancing discriminative cues in high-confidence regions while preserving signal fidelity in low-SNR regions.

To model long-range dependencies, we further introduced Mamba with state-space model parameters Θ = {*A, B, C, D,* Δ}, where *A* ∈ R*^d^*^×*d*^, *B* ∈ R*^d^*, *C* ∈ R*^d^*, *D* ∈ R, and Δ denotes the step size. Discretization of the state-space model produced a one-dimensional causal kernel *k*_Θ_, and the sequence output was computed as:

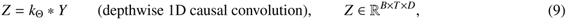

where ∗ denotes channel-wise independent causal convolution. This operation reduced the sequence modeling complexity from *O*(*T* ^2^) to *O*(*Td_s_*) while strengthening both local and long-range interactions through convolutional expansion. In our implementation, the Mamba module used *d*_state_ = 16 and *d*_conv_ = 2. The encoded features were then globally pooled and fed into the classifier to obtain the modification prediction 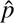 or 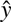. Algorithm 1 summarizes the complete workflow of the proposed model.

**Table 1:**
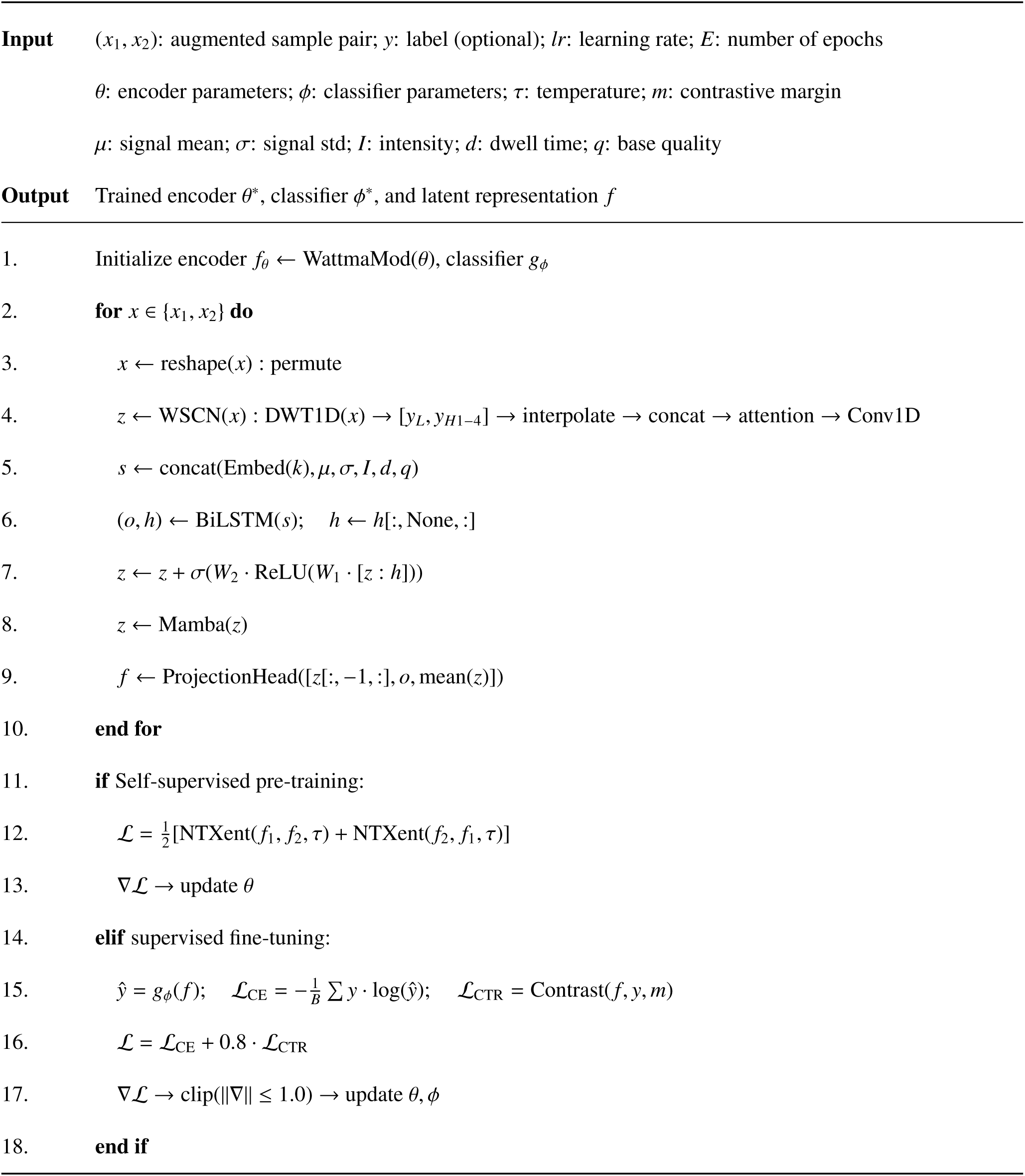
Algorithm 1 — WattmaMod: integrated representation learning framework.

### Model hyperparameter settings and evaluation

During the model training process, we used the AdamW ^53^ optimizer, with an initial learning rate of 0.001 and a batch size of 512. A dynamic learning rate adjustment strategy (ReduceLROnPlateau) was employed, which reduces the learning rate by a factor of 0.1 when the validation metric plateaued for 3 consecutive epochs, with a minimum learning rate of 1×10^−6^. To prevent overfitting, early stopping was enabled to terminate training if no improvement was observed over a certain period. All training was conducted on an NVIDIA A10 GPU. Detailed training hyperparameter settings are summarized in Supplementary Table 3.

We uniformly split each dataset into training and testing sets at a 4:1 ratio. RNA002 undergoes cross-training and validation between IVET, Curlcake (RNA002), and IVT. The RNA004 dataset is based on Curlcake (RNA004) (long sequences) and the rna-modbase in vitro transcription dataset constructed from BED tags (short sequences). The self-supervised pre-training stage ran for 40 epochs, with each epoch lasting approximately 4 minutes to sufficiently capture the distributional characteristics of large-scale data. The supervised fine-tuning was divided into two phases. Phase I: Parameter tuning on 250,000 labeled samples for each modification type, trained for 15 epochs, with each epoch lasting about 2 minutes. Phase II: Partial parameter fine-tuning on 3,000 labeled samples per type, lasting about 3 s per epoch.

Classification performance was evaluated using the true positive rate (TPR, equivalent to recall), false positive rate (FPR), precision, ROC–AUC and PR–AUC. The corresponding definitions were:

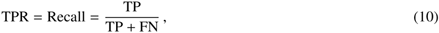

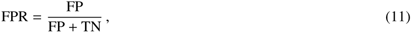

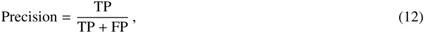

where TP, FP, TN and FN denote the numbers of true positives, false positives, true negatives and false negatives, respectively. Details of the test and analysis datasets are provided in Supplementary Table 2.

## Data availability

Published DRS datasets used in this study were obtained from the work of Novoa lab in the National Center for Biotechnology Information (NCBI) under accession numbers GSE124309^38^ and PRJNA563591^54^, and from the work of Intawat Nookaew under accession number SRP166020^40^. Published *Arabidopsis* and *Populus trichocarpa* DRS datasets are available through ENA and NCBI SRA under accession numbers PRJEB32782^55^ and PRJNA517295^56^, respectively. Published *Saccharomyces cerevisiae* direct RNA sequencing (DRS) datasets are available through ENA under accession number PRJEB37798. Direct RNA sequencing datasets for multiple human cancer cell lines are publicly available in ENA under project accession PRJEB44348. Direct RNA sequencing data of HEK293T wild-type and METTL3-knockout samples are publicly available in ENA under project accession PRJEB40872. The In Vitro Epi-transcriptome (IVET) dataset generated by ONT direct RNA sequencing is publicly available in GEO under accession number GSE227087. Direct RNA sequencing datasets for HeLa wild-type and NSUN2-knockout are publicly available in NCBI under project accession PRJNA872027. RNA004 curlcake synthetic direct RNA sequencing datasets are available from ENA under project accessions PRJEB61874 and PRJEB82528. The RNA004 rna-modbase validation dataset (rna-modbase-validation 2025.03), with defined RNA modifications, is available from the Oxford Nanopore Technologies open-data repository at https://ont-open-data.s3.amazonaws.com/index.html. The processed data generated in this study, including read-level prediction outputs, site-level modification rates, benchmark summary tables, have been deposited in Zenodo: https://doi.org/10.5281/zenodo.18998046. Source data are provided with this paper.

## Code availability

The source code of the WattmaMod is available for research purposes at Github: https://github.com/YuBinLab-QUST/WattmaMod/.

## Acknowledgements

This work was supported by the National Natural Science Foundation of China (No. 62172248), the Natural Science Foundation of Shandong Province of China (No. ZR2021MF098), and the King Abdullah University of Science and Technology (KAUST) Office of Research Administration (ORA) under Award No REI/1/5234-01-01, REI/1/5414-01-01, REI/1/5289-01-01, REI/1/5404-01-01, REI/1/5992-01-01, URF/1/4663-01-01.

## Author contributions

R.H. conceived the study, and provided project supervision and resource support. B.Y. and X.S. jointly designed and developed the research framework, and made the major intellectual contributions to the work. X.S. carried out the principal data processing, model development, experimental implementation, performance evaluation, and drafted the manuscript. X.L., J.Q., and T.Y. provided assistance in data preparation and partial experimental validation. X.G. contributed to guidance on manuscript preparation. All authors reviewed and approved the final manuscript.

## Competing interests

The authors declare no competing interests.

## Supplementary information

Supplementary information is available in the online version of the paper.

## Notes

### Competing Interest Statement

The authors have declared no competing interest.

