## Supplemental figure for "WattmaMod enables high-resolution and extensible RNA modification profiling for nanopore direct RNA sequencing"

### Supplementary notes

#### Supplementary Note 1: Model design and feature characterization

Supplementary Figs. S1–S3 provide additional evidence on model architecture and feature patterns underlying modified base recognition. Supplementary Fig. S1 outlines the main modules. Supplementary Fig. S2 shows event-level feature distributions and feature contributions. Supplementary Fig. S3 summarizes sequence context enrichment and low-dimensional event representations across modification types.

[†]These authors contributed equally.

#### Supplementary Note 2: Robustness, generalization and operating behaviour

Supplementary Figs. S4–S9 and Fig. S12 evaluate model robustness, generalization, and operating behaviour across datasets and conditions. Supplementary Fig. S4 examines the effects of training datasets and k-mer resolution on cross-dataset performance. Supplementary Fig. S5 assesses ROC/PR behaviour under different confidence strategies and read-length settings. Supplementary Fig. S6 visualizes representations learned in multi-modification tasks with baseline comparisons. Supplementary Fig. S7 summarizes the data augmentation design, and Supplementary Fig. S8 shows enrichment performance by lift curves. Supplementary Fig. S9 presents site-level analysis of synthetic Curlcake RNA004 data, highlighting pattern- and coverage-dependent trends. Supplementary Fig. S12 demonstrates multi-modification evaluation and feature space visualization on RNA004 oligonucleotide/IVT benchmark data, providing validation for short reads.

#### Supplementary Note 3: Biological validation and downstream analyses

Supplementary Figs. S10–S11 and S13–S21 provide biological validation and downstream analyses across multiple species, cell types, and perturbation settings. Supplementary Figs. S10 and S11 show the positional distribution and functional/pathway enrichment of high-confidence DRACH sites in HEK293T cells. Supplementary Figs. S13 and S14 examine modification changes under knockout or mutant conditions. Supplementary Figs. S15–S17 summarize co-occurrence, motif preference, and structure-related patterns. Supplementary Figs. S18–S21 further present cross-batch reproducibility, the effect of fine-tuning set size, sequence-context signatures across modifications, and the gene-level distribution of predicted $m^6A$ sites.

### Supplementary methods

To ensure consistent and reproducible preprocessing across different nanopore chemistries, we implemented two standard signal-to-reference mapping workflows for RNA002 and RNA004 datasets (Table S1). Table S2 provides an overview of the datasets used in this study. For RNA002, raw FAST5 files were basecalled using Guppy (v6.1.5) to generate read sequences and per-base information, followed by sequence alignment with minimap2 (v2.22; `-ax map-ont`). We then applied Tombo resquiggle (v1.5.1; `-fit-global-scale -include-event-stdev`) to obtain signal-to-reference mappings and event-level statistics for downstream feature extraction. For RNA004, raw reads were processed in POD5 format using the pod5 toolkit (v0.3.35) when necessary (FAST5→POD5 conversion), and basecalling/alignment were performed with Dorado (v0.9.6) using RNA004-specific models with `-emit-moves` to retain move information. Signal-level alignment to the reference was subsequently computed using Uncalled4 (v4.1.0; `-rna -norm-iterations 1`), producing refined signal-to-reference coordinates compatible with our training and evaluation pipeline. Table S3 summarizes the software versions and key non-default options used in this study. Training configuration and hyperparameters are provided in Supplementary Section .

**Table S1:** Software tools and key non-default options used for basecalling and signal-to-reference mapping.

| Chemistry | Step | Version | Key non-default options / notes |
|---|---|---|---|
| RNA002 | Basecalling | Guppy | v6.1.5; `-recursive`; optional FAST5 output: `-fast5_out` |
| RNA002 | FAST5 conversion | ont_fast5_api | v4.0.0; `multi_to_single_fast5 -recursive` |
| RNA002 | Alignment (sequence) | minimap2 | v2.22; `-ax map-ont` |
| RNA002 | Signal-to-reference mapping | Tombo (resquiggle) | v1.5.1; `-overwrite -fit-global-scale -include-event-stdev` |
| RNA004 | Signal container conversion | pod5 | v0.3.35; convert FAST5 to POD5 for downstream processing (`pod5 convert fast5`) |
| RNA004 | Basecalling | Dorado | v0.9.6; models `rna004_130bps_sup@v5.1.0` or `rna004_70bps_hac@v5.1.0`; `-emit-moves -emit-sam` |
| RNA004 | Alignment (sequence) | Dorado / minimap2 | BAM produced during Dorado basecalling; alignment configured for ONT RNA reads (e.g., `-x map-ont` equivalent) |
| RNA004 | Signal-to-reference mapping | Uncalled4 | v4.1.0; `-ref REF.fa -rna -norm-iterations 1` |

**Note:** RNA002 datasets were processed with a Guppy→minimap2→Tombo workflow, whereas RNA004 datasets were processed with a Dorado→Uncalled4 workflow to obtain signal-to-reference mappings compatible with downstream feature extraction.

**Table S2:** Dataset sources and summary

| Dataset | Chemistry | Modification / control | Usage |
|---|---|---|---|
| IVET (in vitro + enzymatic treatment) | RNA002 | $m^6A$, $m^5C$, $m^1A$; canonical A/C controls | FT |
| IVT (in vitro transcription, multi-mod set) | RNA004 | $m^6A$, $m^5C$, $m^1A$, $hm^5C$, inosine, Ψ, $m^7G$, f5C; canonical controls (A/C/G/U) | FT |
| Curlcake (synthetic constructs) | RNA002 | $m^6A$, $m^5C$, $m^1Ψ$, $m^5U$; unmodified control | FT |
| Curlcake (synthetic constructs) | RNA004 | $ac^4C$, $hm^5C$, $m^1Ψ$, $m^5C$, $m^6A$, $m^5U$, Ψ; unmodified control | FT |
| RNA-modbase (synthetic oligo benchmark) | RNA004 | $m^6A$, $m^5C$, Ψ, inosine; unmodified control | FT / Val |
| HEK293T (WT vs METTL3 KO) | RNA002 | $m^6A$, $m^5C$, $m^1A$, $hm^5C$, inosine, Ψ, $m^7G$ | Pretrain/ Val / Downstream |
| HEK293T native DRS | RNA004 | $m^6A$-focused analysis | Val / Downstream |
| A549 / HCT116 / MCF-7 / HepG2 / K562 / HEYA8 / H9 | RNA002 | $m^6A$-centered analysis with additional nine modification types | Downstream |
| HeLa (WT vs NSUN2 KO) | RNA002 | $m^5C$-focused analysis | Downstream |
| *Arabidopsis thaliana* (Col-0 / vir-1 / virc) | RNA002 | $m^6A$-focused analysis | Downstream |
| *Populus trichocarpa* | RNA002 | $m^6A$, $m^1A$ | Downstream |
| *S. cerevisiae* (WT vs methyltransferase-deficient KO; rRNA) | RNA002 | $m^6A$ (rRNA) | Downstream |

**Definitions:** IVT = in vitro transcription with predefined modified vs canonical controls; IVET = in vitro transcription with enzymatic treatment and matched control design; Curlcake/RNA-modbase = synthetic constructs/oligos with predefined modification sites. **Notation:** canonical RNA bases are A/C/G/U (not T); Ψ denotes pseudouridine; $m^1Ψ$ denotes 1-methylpseudouridine. **Usage:** Pretrain = self-supervised pre-training; FT = supervised fine-tuning with labeled sites/reads; Val = validation only; Downstream = biological profiling/association analyses (typically weak-label setting).

### Supplementary figures

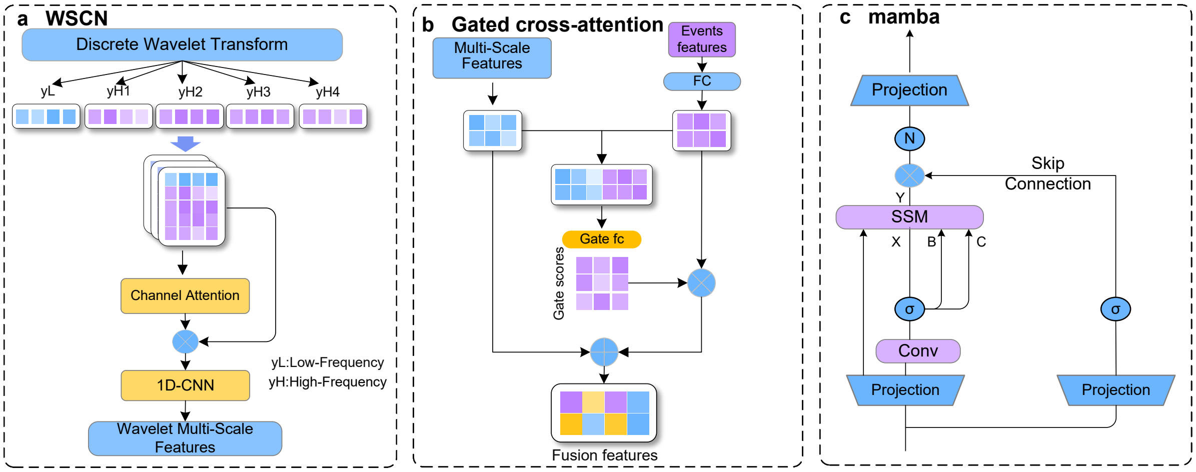


**Figure S1:** Schematic overview of WattmaMod. **a**, Architecture of the wavelet-guided signal encoder, including discrete wavelet decomposition, channel-wise reweighting and one-dimensional convolution blocks. **b**, Gated cross-attention fusion of wavelet-derived signal features and event-level descriptors. **c**, Mamba-based sequence encoder for long-range context modeling.

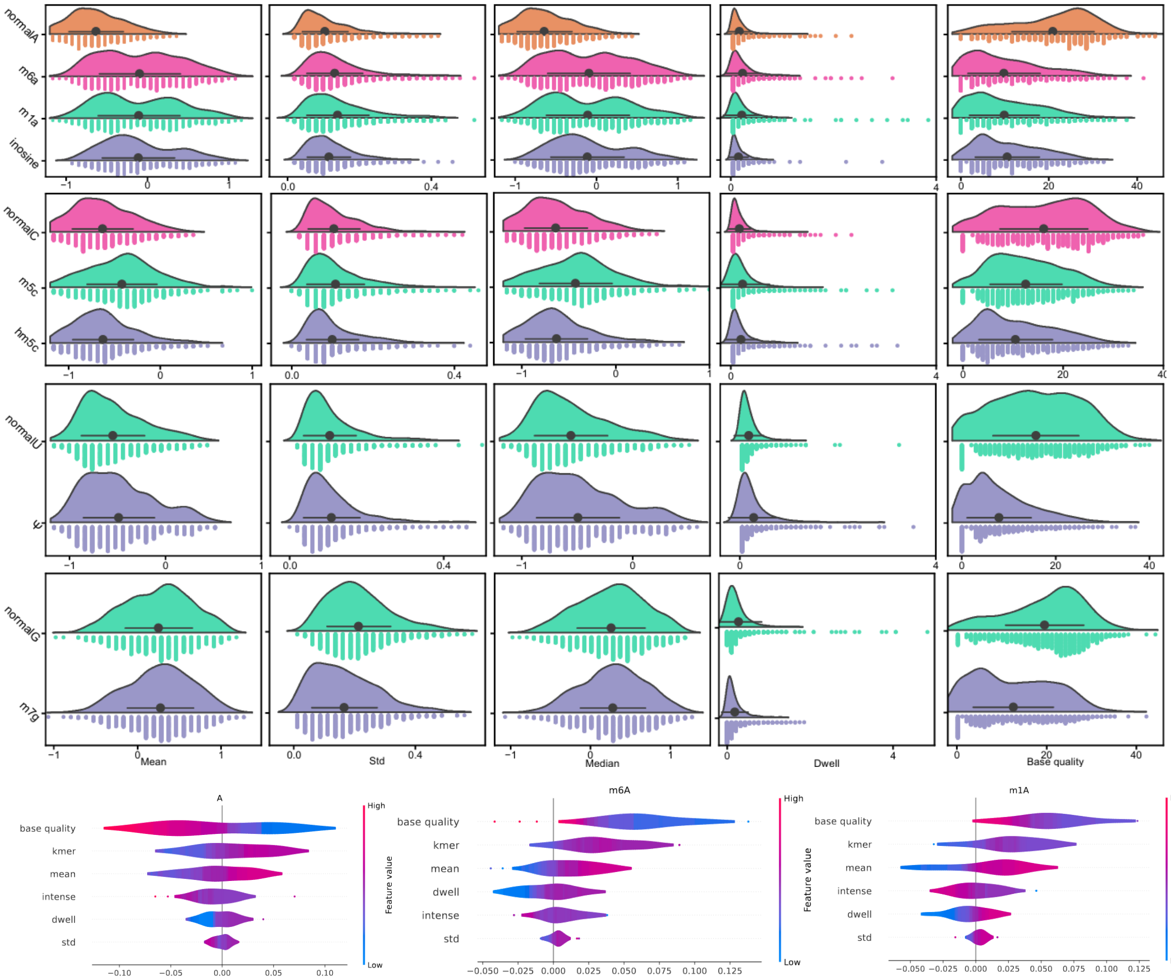


**Figure S2:** Event-level feature distributions and feature attribution for canonical and modified nucleotides. **a**, Raincloud plots of five event-level descriptors, including mean current, standard deviation, median, dwell time and base quality, for canonical bases (A/C/G/U) and their corresponding modification classes ($m^6A$, $m^1A$, inosine, $m^5C$, $hm^5C$, Ψ and $m^7G$). Points represent event observations extracted from the target-centered window, and violin shapes indicate the distribution of each feature. **b**, SHAP value distributions of major input descriptors, including k-mer context, base quality and event statistics, for discrimination among A, $m^6A$ and $m^1A$. Larger absolute SHAP values indicate stronger contributions to model prediction.

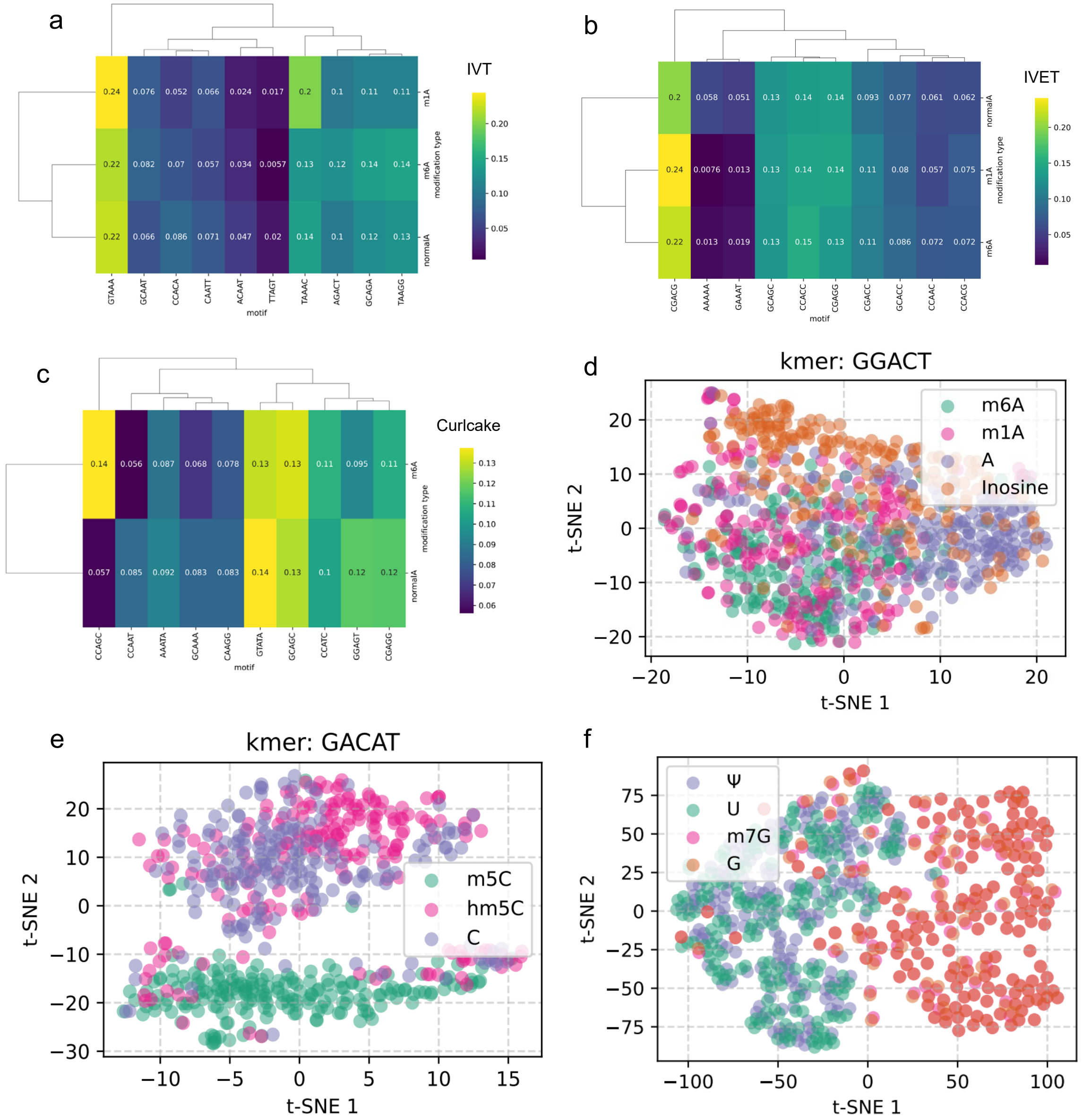


**Figure S3:** Sequence-context enrichment and low-dimensional visualization of event features across RNA modification types. **a–c**, Heatmaps of normalized 5-mer enrichment for modification classes relative to controls in the IVT (**a**), IVET (**b**) and Curlcake (**c**) datasets. Enrichment was calculated from target-centered 5-mer contexts of labeled sites, and both rows and columns were hierarchically clustered to highlight shared and modification-specific sequence preferences. **d–f**, t-SNE projections of event-level feature vectors for representative sequence contexts: GGACT for A/$m^6$A/$m^1$A/inosine (**d**), GACAT for C/$m^5$C/$hm^5$C (**e**) and U/Ψ/$m^7$G/G (**f**). Each point represents one aligned event, and colours indicate the ground-truth class.

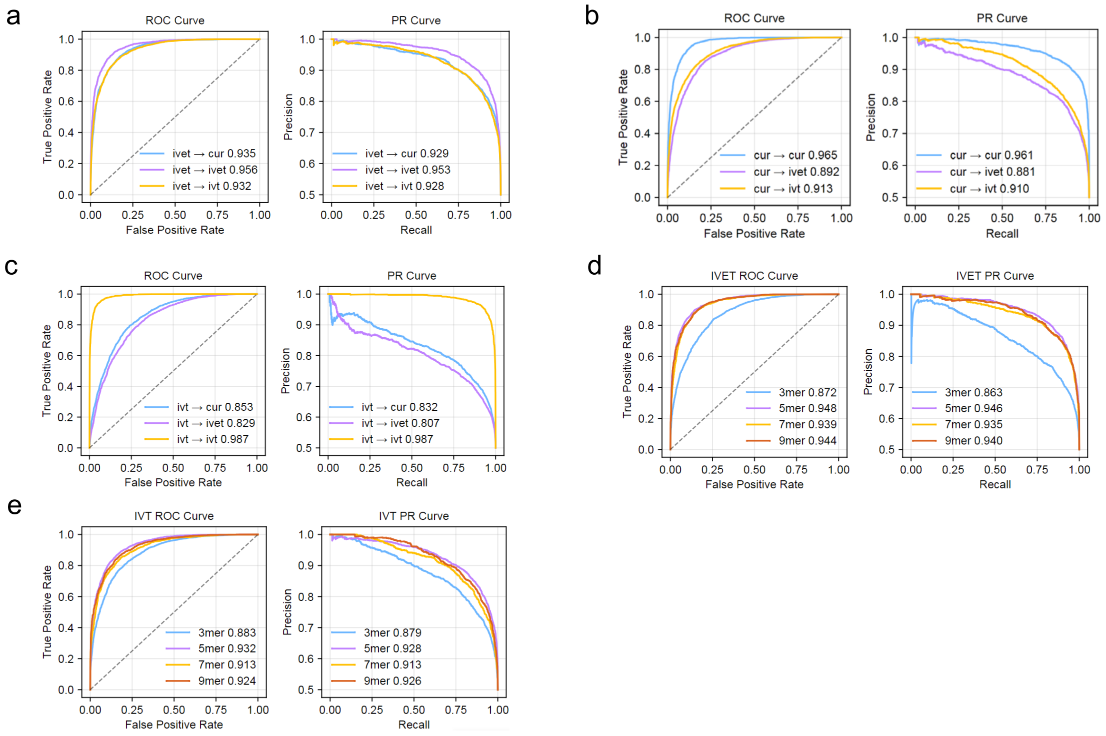


**Figure S4:** Model performance across training datasets and k-mer lengths. **a**, ROC (left) and precision–recall (PR; right) curves for a model trained on IVET and evaluated on the IVET test set and Curlcake. **b**, ROC and PR curves for a model trained on Curlcake and evaluated on Curlcake and the IVET test set. **c**, ROC and PR curves for a model trained on IVT and evaluated on the IVT test set and Curlcake. **d**, Effect of k-mer length (3-, 5-, 7- and 9-mer) on performance for models trained on IVET and evaluated on the IVET test set. **e**, Cross-dataset generalization to IVT for models trained on IVET with different k-mer lengths and evaluated on the IVT test set. **f**, Cross-dataset generalization to IVET for models trained on IVT with different k-mer lengths and evaluated on the IVET test set.

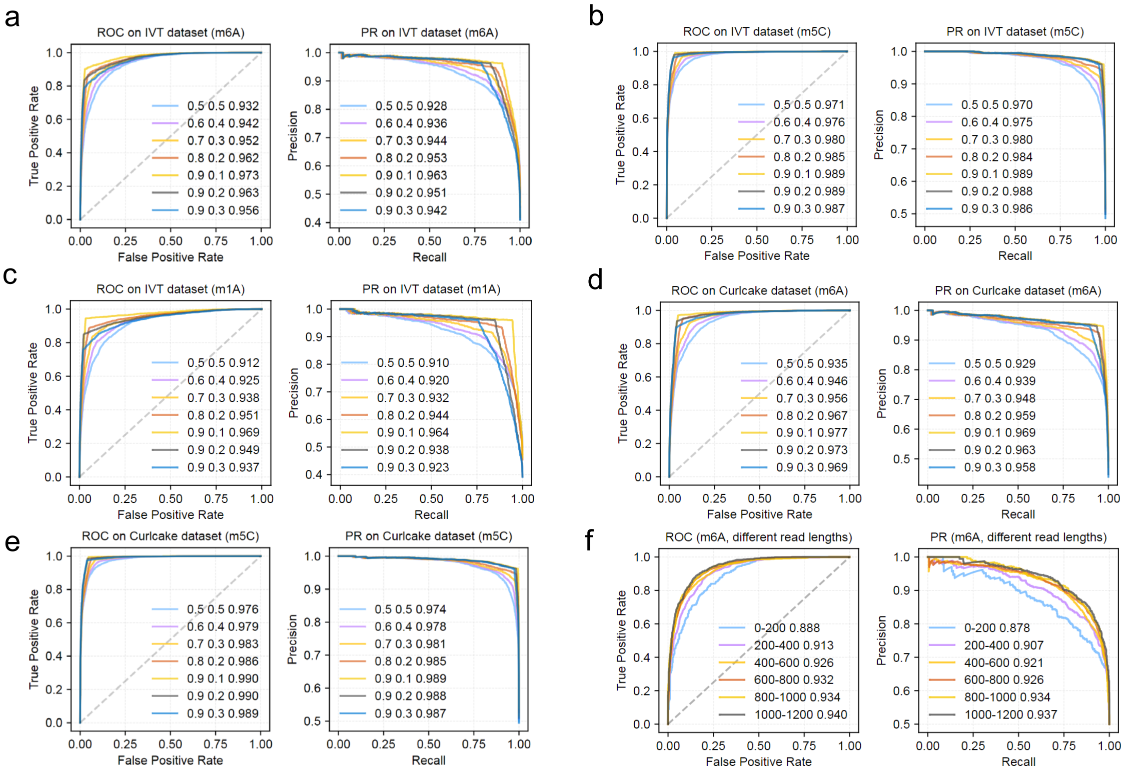


**Figure S5:** Sensitivity of ROC and PR performance to dual-threshold confidence strategies. **a–c**, ROC and PR curves on IVT for $m^6A$ (**a**), $m^5C$ (**b**) and $m^1A$ (**c**) under multiple ($t_{pos}, t_{neg}$) settings, where high-confidence positives and negatives were defined by the two thresholds. **d**, ROC and PR curves for $m^6A$ on Curlcake evaluated using the same set of threshold pairs. **e**, ROC and PR curves for $m^5C$ on Curlcake under the corresponding confidence strategies. **f**, ROC and PR curves for $m^6A$ stratified by read-length bins (0–200, 200–400, 400–600, 600–800, 800–1000 and 1000–1200 nt), showing performance across length-defined subsets.

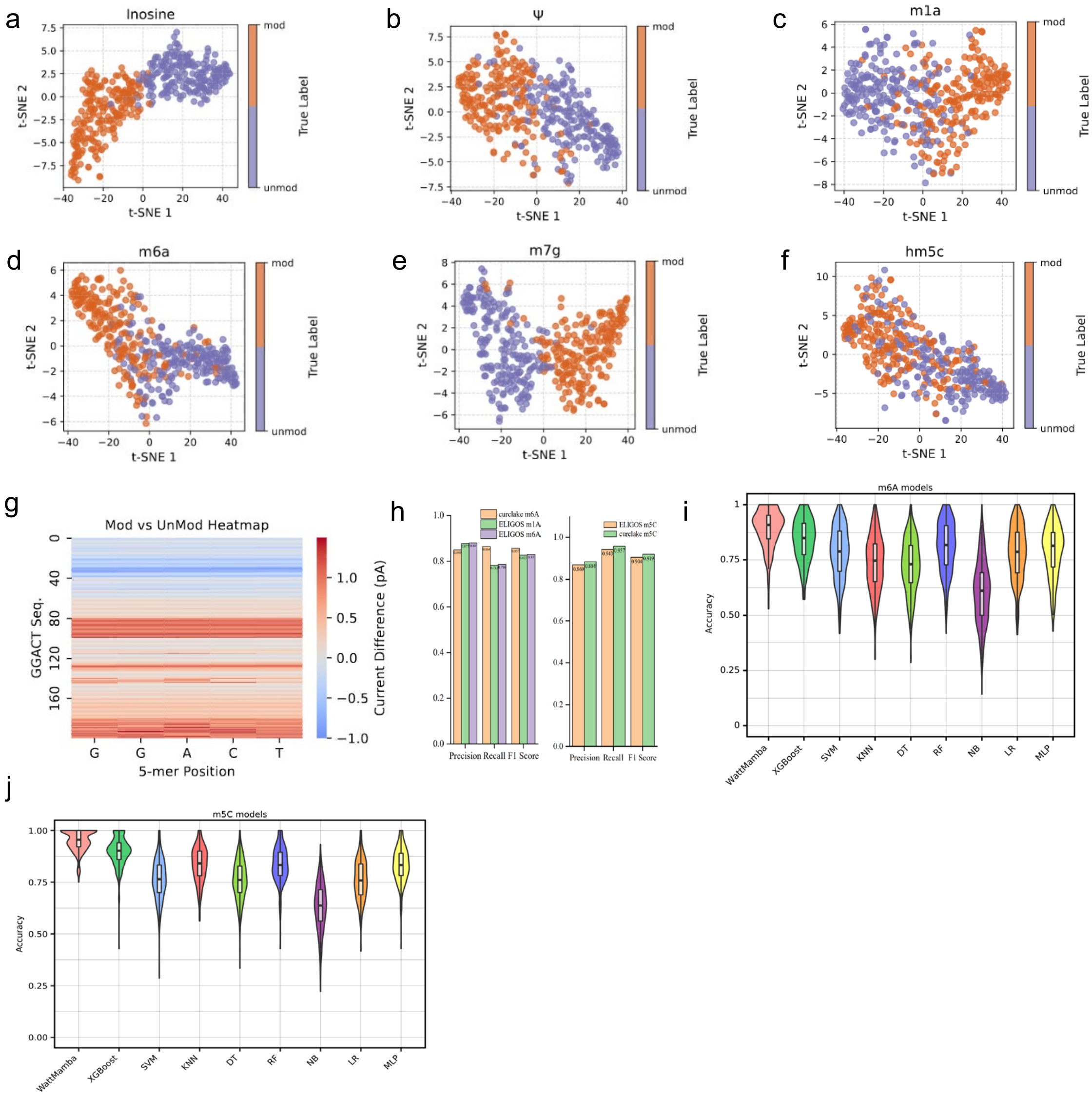


**Figure S6:** Representation visualization, motif-level signal differences and model comparisons. **a–f**, t-SNE projections of learned embeddings after supervised fine-tuning for inosine (**a**), Ψ (**b**), $m^1A$ (**c**), $m^6A$ (**d**), $m^7G$ (**e**) and $hm^5C$ (**f**); each point denotes one target-centered event or window and is coloured by the ground-truth label. **g**, Heatmap of normalized current differences between modified and unmodified groups for the GGACT motif, highlighting position-dependent shifts around the target base. **h**, Precision, recall and F1-score comparisons on the Curlcake and IVET test sets for representative modification tasks. **i**, Classification accuracy comparison for the $m^6A$ task between WattmaMod and classical baseline models, including XGBoost, SVM, KNN, decision tree, random forest, Naive Bayes, logistic regression and MLP. **j**, Classification accuracy comparison for the $m^5C$ task using the same set of baseline models.

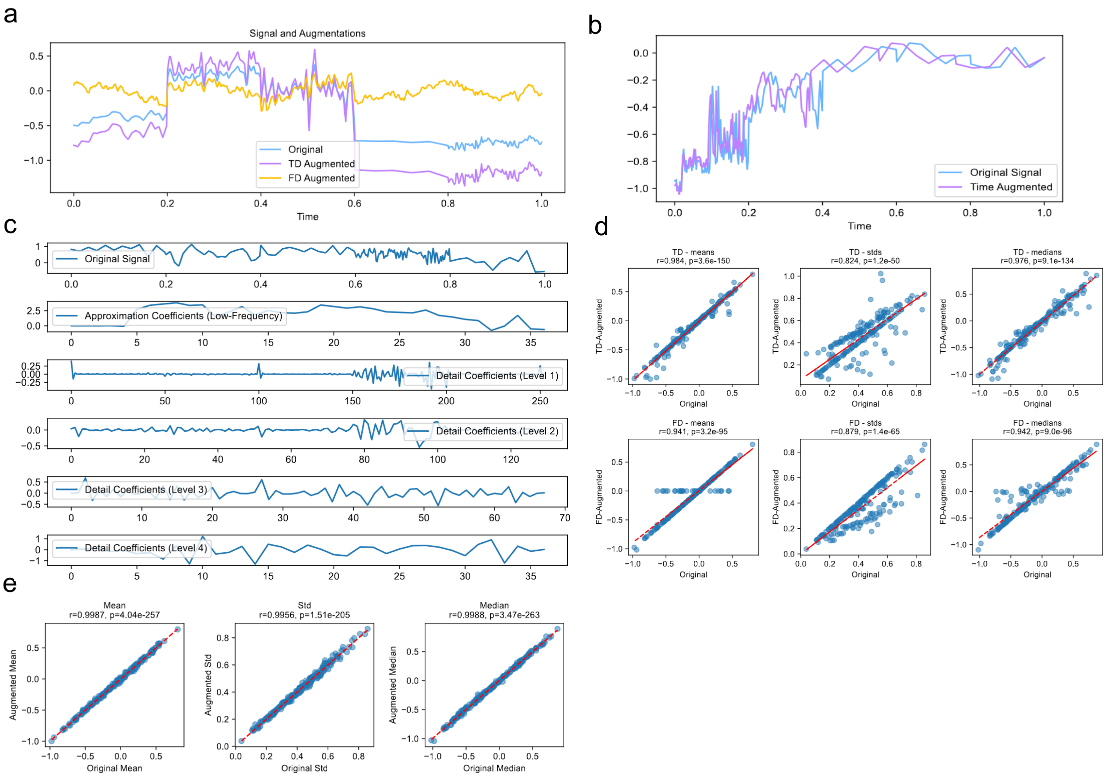


**Figure S7:** Evaluation of signal augmentation strategies and wavelet-derived transformations. **a**, Representative raw current traces overlaid with augmented signals generated by time-domain augmentation (TDA) and frequency-domain augmentation (FDA). **b**, Comparison of original and augmented signal examples, illustrating diverse perturbation patterns while preserving the overall signal structure. **c**, Four-level discrete wavelet transform (DWT) decomposition of a representative signal into one approximation component and detail components at levels 1–4. **d**, Correlations between augmented and original signals measured by summary statistics, including the mean, standard deviation and median, for TDA (top) and FDA (bottom). **e**, Correlation plots showing how supervised augmentations preserved statistical consistency with the original signals across the same three descriptors.

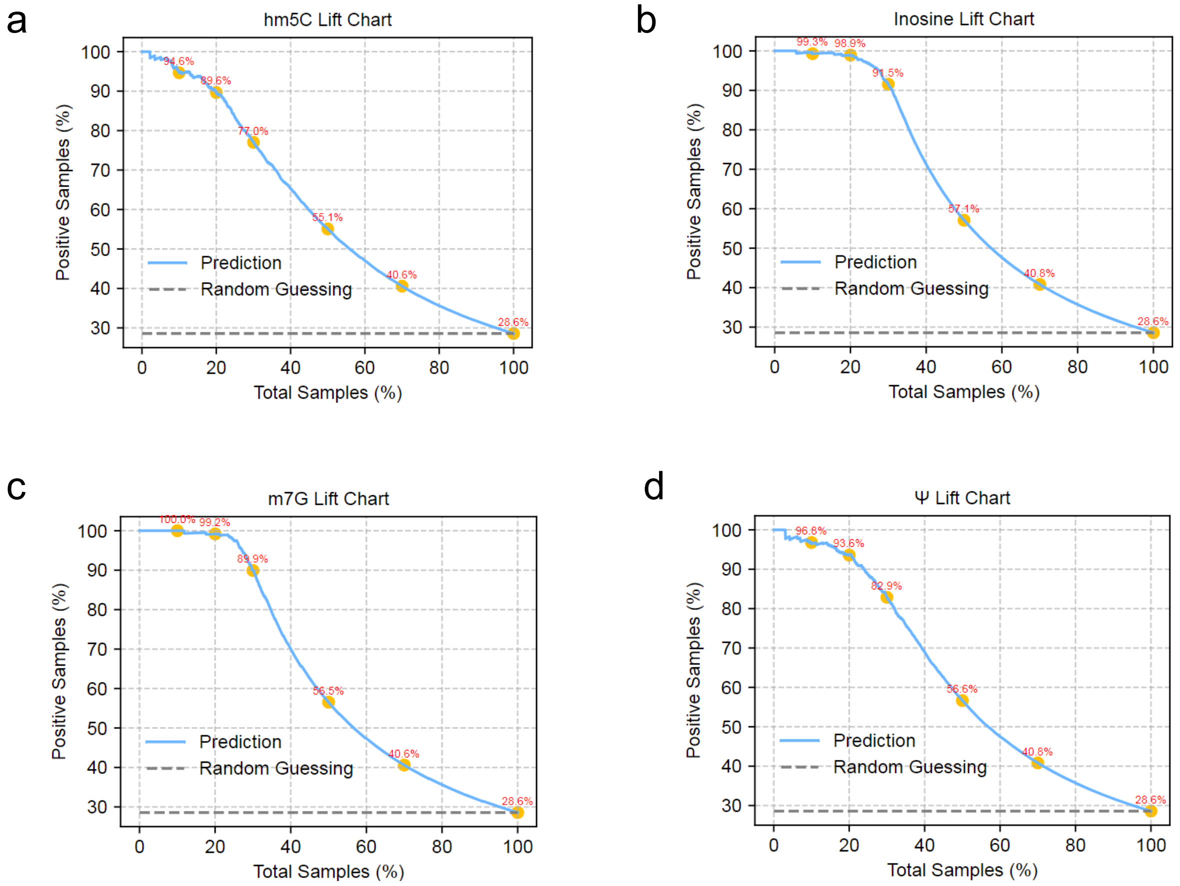


**Figure S8:** Lift-curve analysis for multi-modification detection. **a–d**, Lift curves for $hm^5C$ (**a**), inosine (**b**), $m^7G$ (**c**) and Ψ (**d**). Samples were ranked by predicted modification probability and accumulated along the x axis as the sampling proportion, whereas the y axis indicates the cumulative fraction of true positives recovered. Dashed lines indicate the random-ranking baseline.

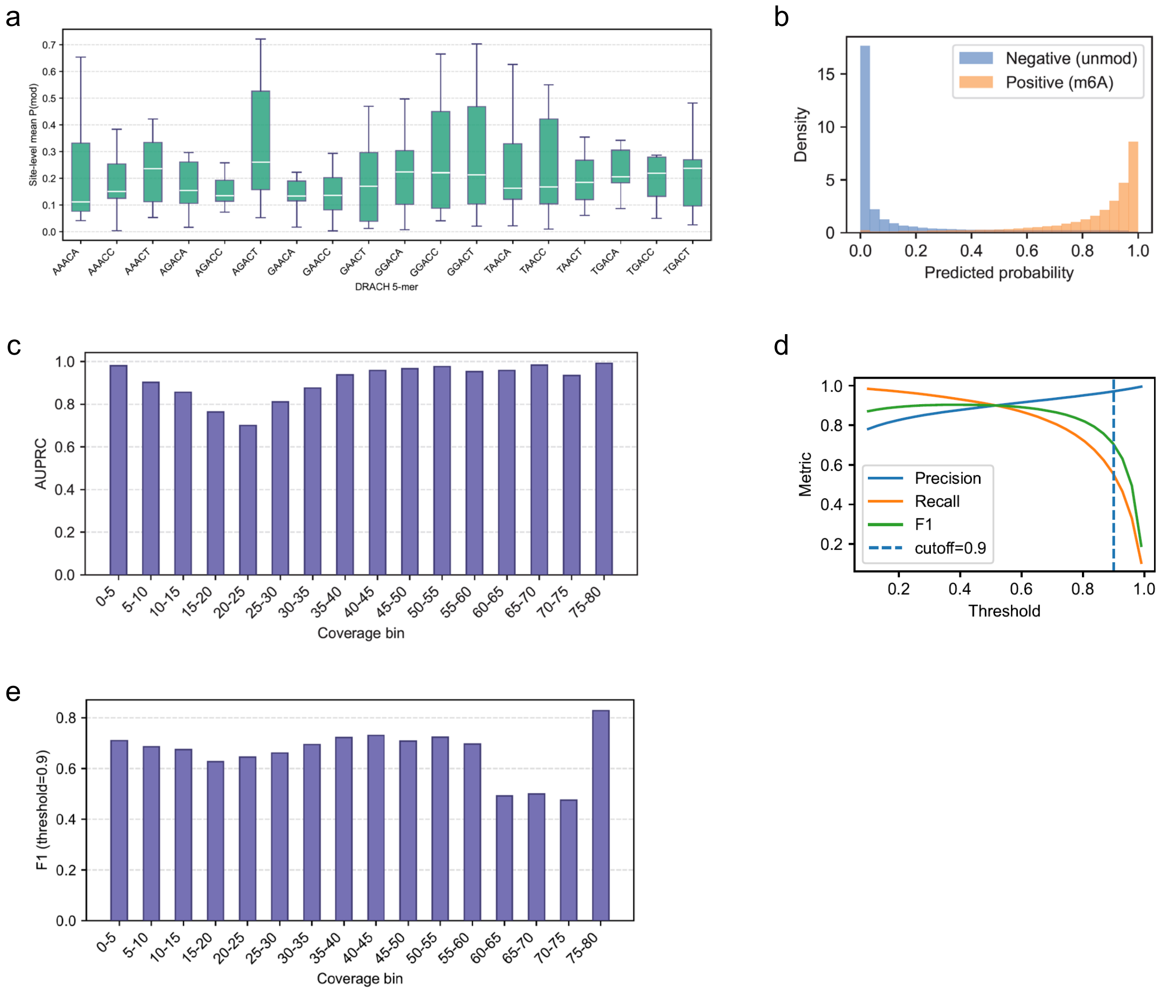


**Figure S9:** Single-molecule-level evaluation of $m^6A$ detection on the RNA004 synthetic Curlcake dataset. **a**, Boxplots of site-level mean predicted probabilities across DRACH 5-mer contexts, summarizing motif-dependent variation in predicted $m^6A$ levels. **b**, Density distributions of predicted probabilities for unmodified controls and $m^6A$-modified sites. **c**, Site-level AUPRC stratified by read-coverage bins, showing the dependence of performance on coverage. **d**, Precision, recall and F1 score as functions of the decision threshold, with the selected operating point indicated. **e**, F1 score across coverage bins at the selected operating threshold, highlighting the coverage range in which site-level calls were most reliable.

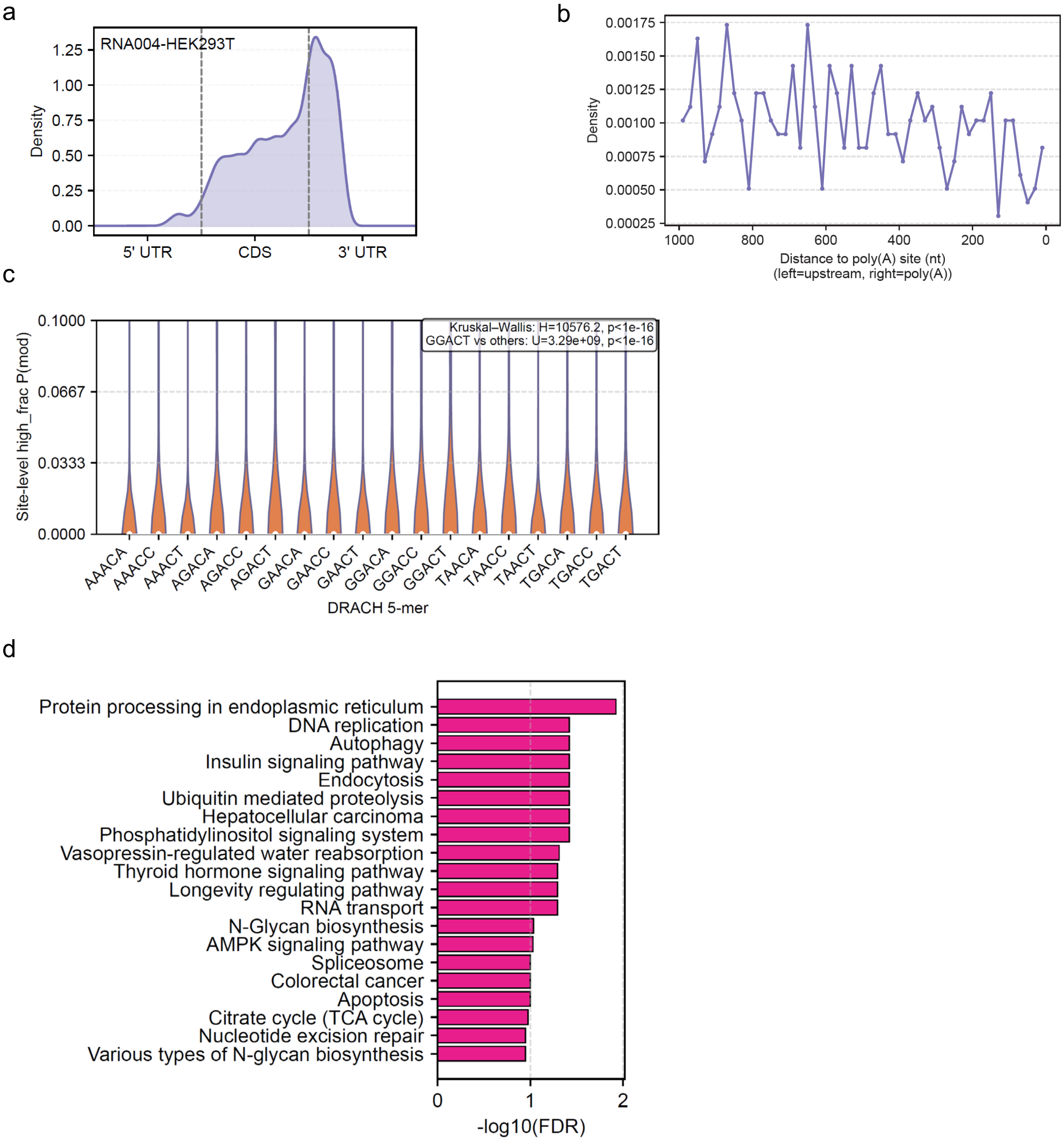


**Figure S10:** Transcript positional distribution and functional enrichment of high-confidence DRACH sites in HEK293T cells generated using RNA004. **a**, Metagene profiles of DRACH sites mapped onto rescaled 5′UTR, CDS and 3′UTR coordinates, showing their relative enrichment along transcripts. **b**, Poly(A)-anchored profiles of DRACH sites aligned to transcript 3′ ends, summarizing the distance-dependent distribution upstream of cleavage sites. **c**, Distributions of site-level high-confidence read fractions across 18 DRACH 5-nt motifs, highlighting motif-dependent variability. **d**, Gene Ontology biological process enrichment analysis for genes harboring high-confidence sites, showing the top-ranked terms by adjusted significance.

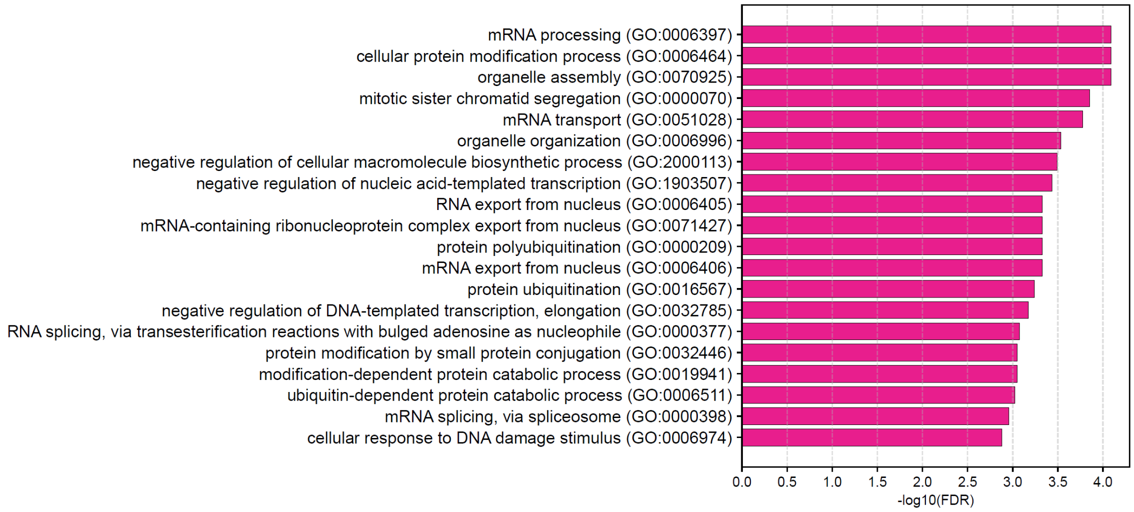


**Figure S11:** Pathway enrichment of genes harboring high-confidence DRACH sites in HEK293T cells generated using RNA004. KEGG pathway enrichment analysis showing the top enriched pathways ranked by adjusted significance.

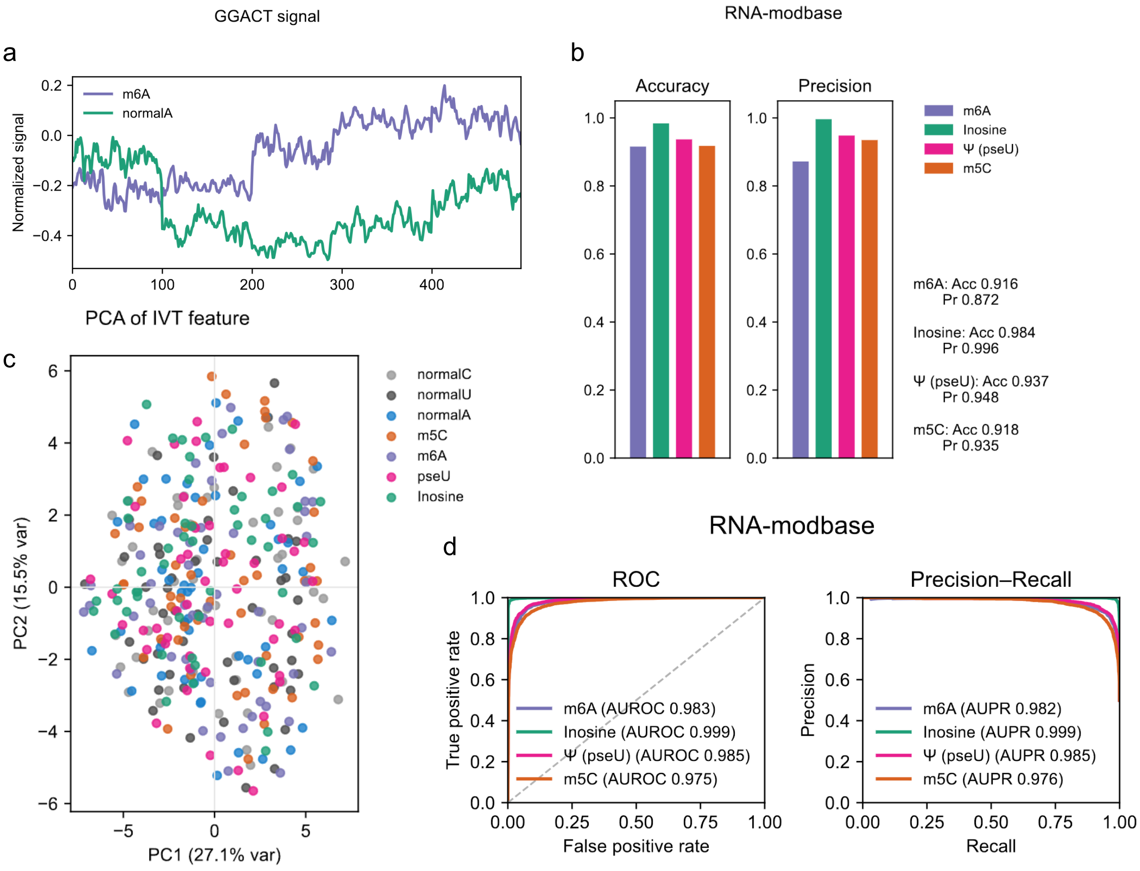


**Figure S12:** Multi-modification detection on the RNA004 in vitro transcription dataset (RNA-modbase) and feature-space visualization. **a**, Example motif-level signal comparison for GGACT, showing mean normalized current traces for modified and unmodified groups across the target-centered window. **b**, Summary of overall read-level performance across $m^6A$, inosine, Ψ and $m^5C$ tasks, reported as accuracy and precision. **c**, PCA projection of RNA004 in vitro transcription feature windows from multiple classes, including canonical controls and modification types, illustrating class separability in the learned feature space. **d**, ROC and precision–recall curves for the four modification types on the same dataset.

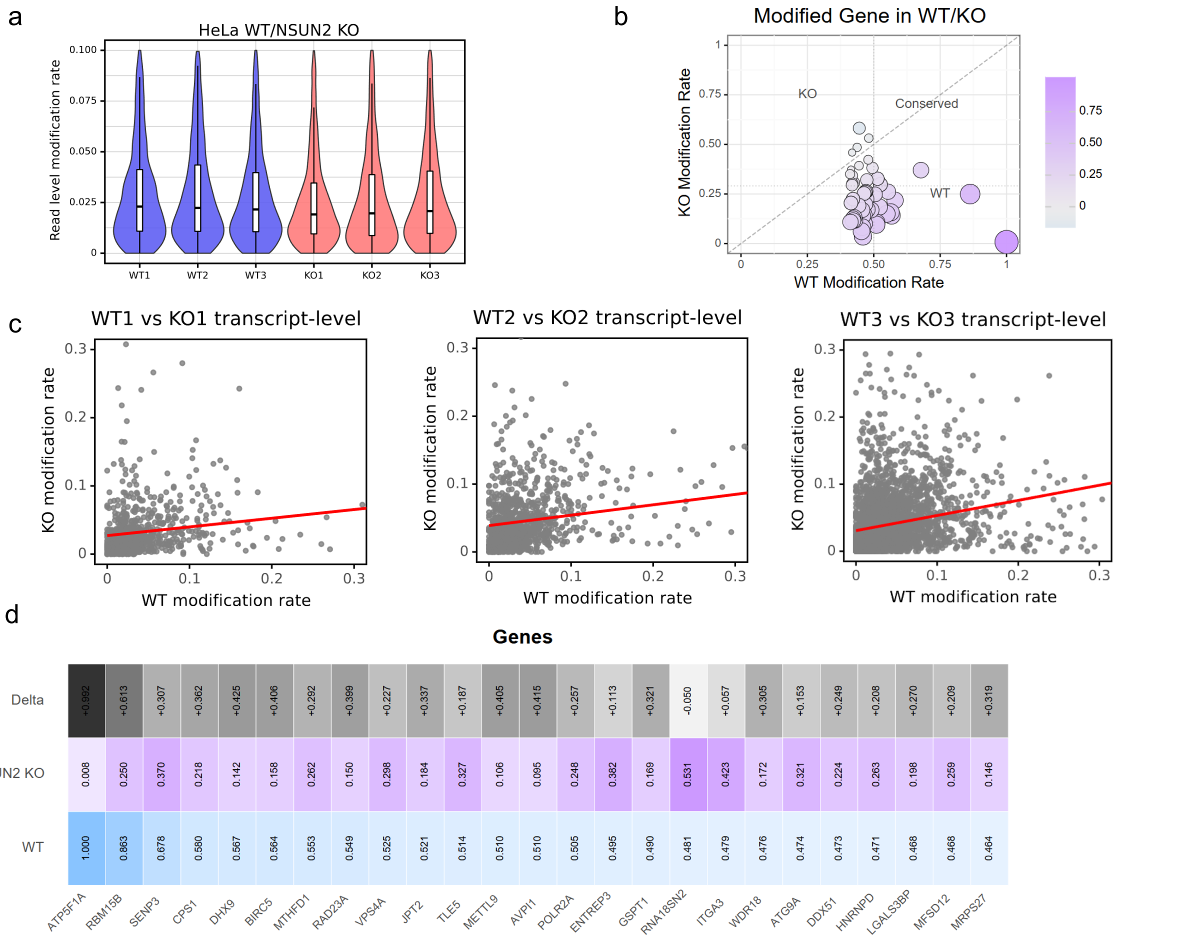


**Figure S13:** $m^5C$ landscape in wild-type and NSUN2-knockout HeLa cells. **a**, Distributions of read-level $m^5C$ modification rates across biological replicates from wild-type (WT) and NSUN2-knockout (KO) samples. **b**, Bubble plot of the top transcripts with high $m^5C$ levels in WT and their corresponding changes in KO; bubble size indicates the magnitude of the rate difference, and colour denotes the direction of change. **c**, Pairwise transcript-level comparisons between WT and KO replicates, showing concordance across samples and conditions. **d**, Heatmap of transcript-level $m^5C$ rates in WT and KO together with the corresponding changes for representative $m^5C$-enriched genes.

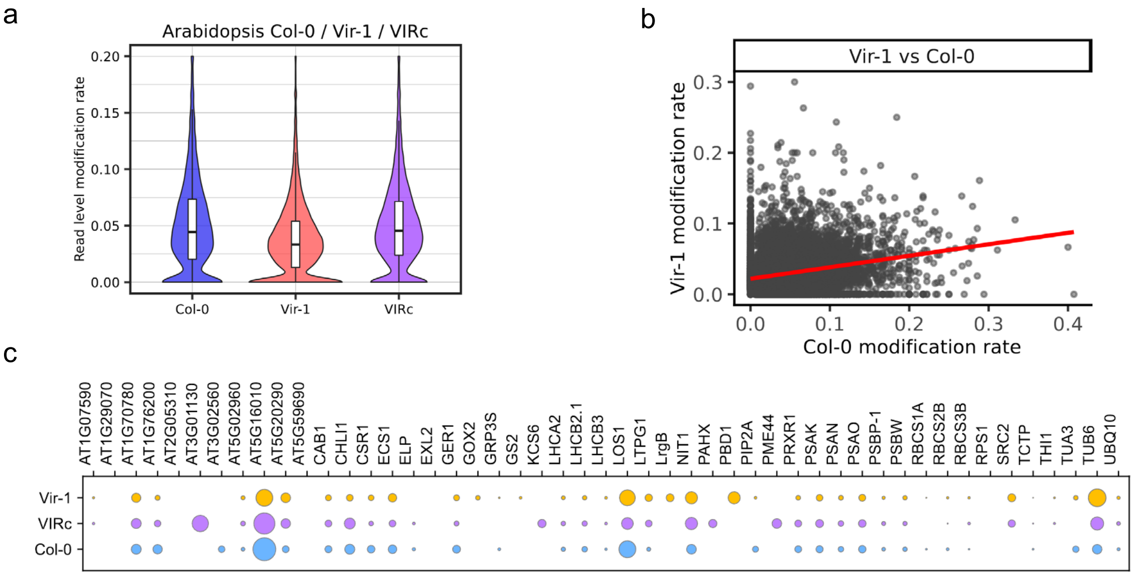


**Figure S14:** m$^6$A patterns in *Arabidopsis thaliana* wild-type and VIR-mutant samples. **a**, Distributions of read-level m$^6$A modification rates in Col-0, vir-1 and vir-c samples. **b**, Transcript-level comparison of m$^6$A rates between vir-1 and Col-0, summarizing global concordance and overall shifts. **c**, Bubble plot summarizing gene-wise m$^6$A rates across Col-0, vir-1 and vir-c, enabling direct comparison of modification levels across genotypes.

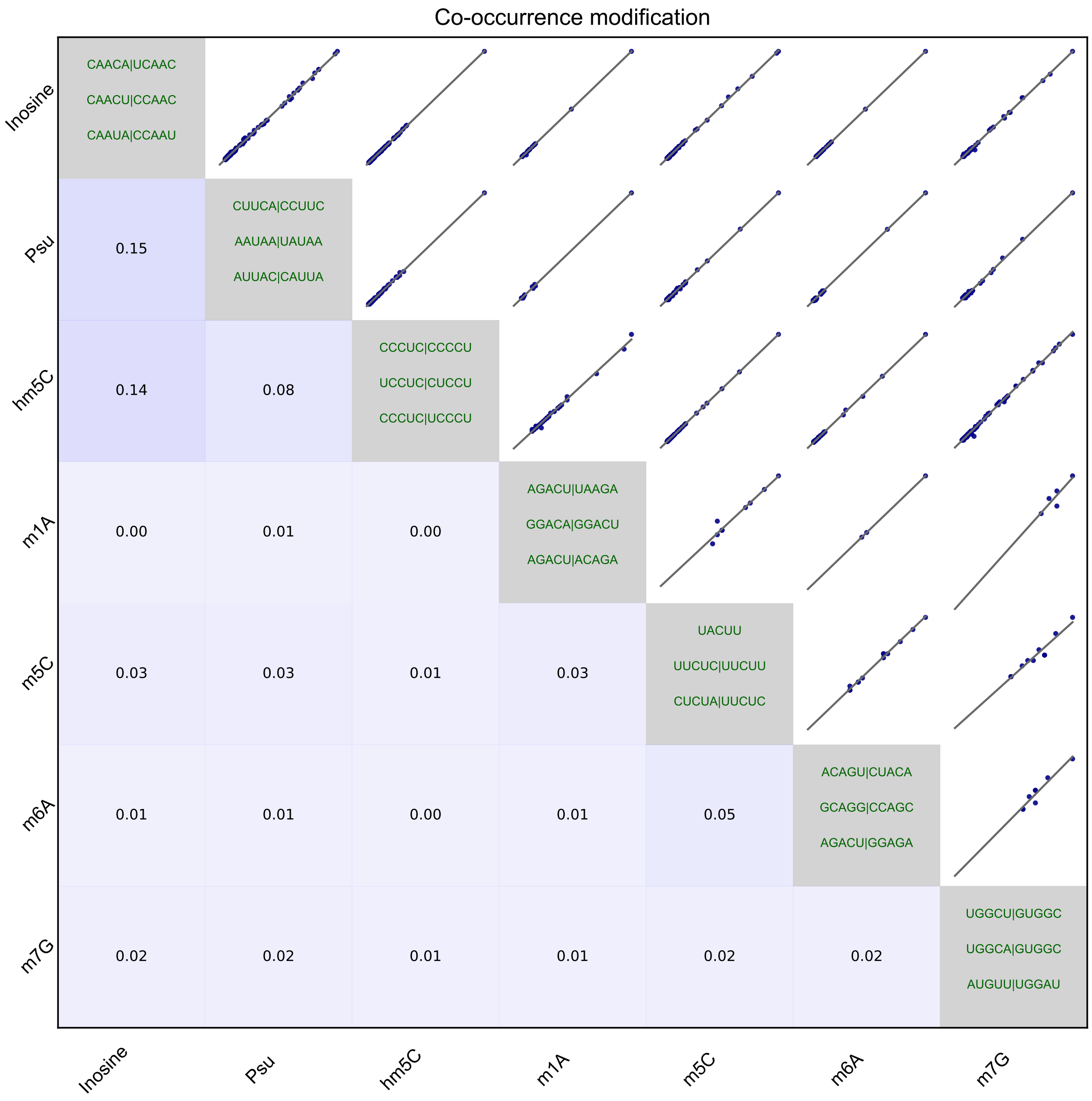


**Figure S15:** Predicted co-occurrence of RNA modifications in human cells. Heatmap of pairwise co-occurrence among seven RNA modification types, including $m^6A$, $m^5C$, $hm^5C$, $m^1A$, Ψ, $m^7G$ and inosine, within a local window around predicted sites. The lower triangle shows Jaccard similarity between modification-specific site sets, the upper triangle shows coverage-dependent associations, and the diagonal displays representative enriched motifs for each modification type.

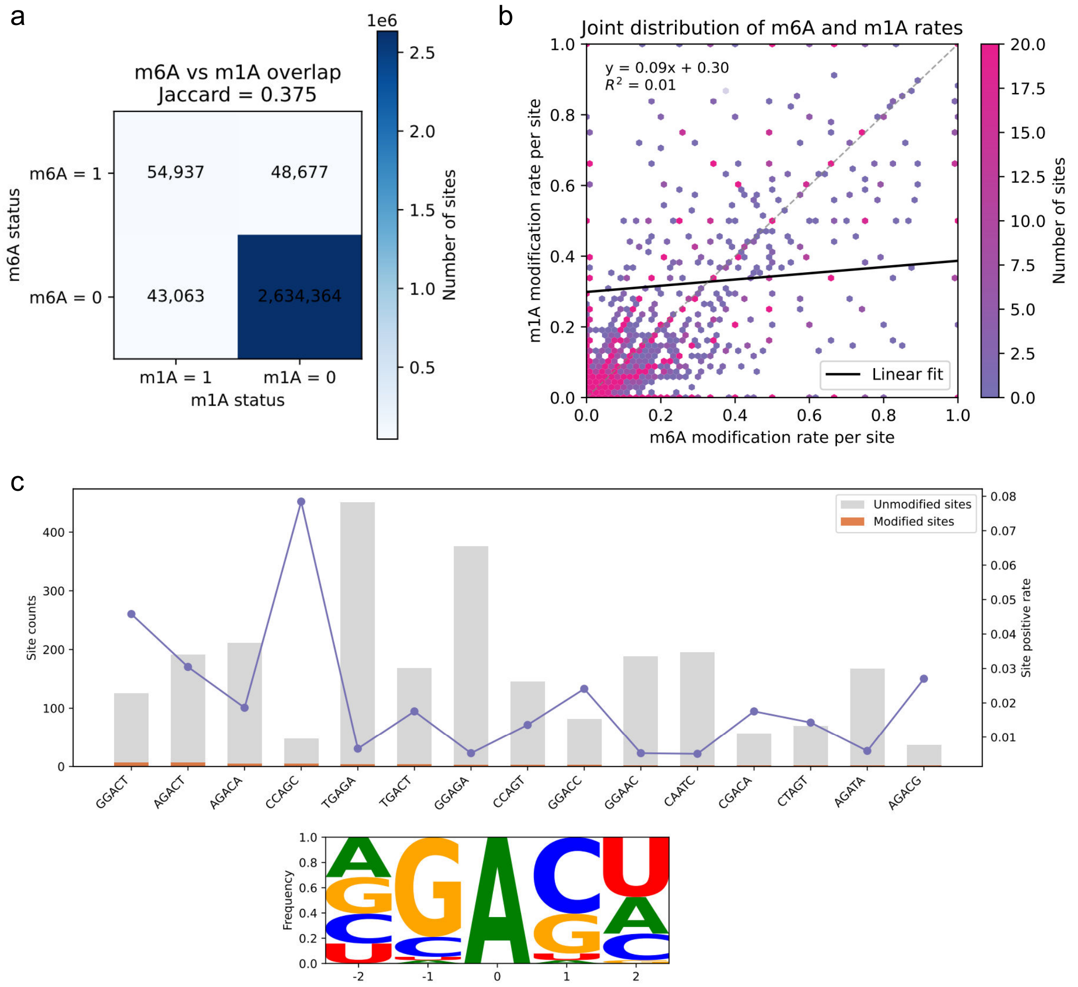


**Figure S16:** Motif preferences and site-level co-modification of $m^6A$ and $m^1A$ in the *Populus trichocarpa* transcriptome. **a**, Site-level co-modification map of $m^6A$ and $m^1A$ at single-nucleotide resolution, summarizing overlapping and modification-specific sites. **b**, Joint distribution of site-level modification rates for $m^6A$ and $m^1A$, illustrating concordance and divergence across sites. **c**, Sequence-context preferences of $m^6A$ sites, showing the top enriched A-centered 5-mers and the corresponding sequence logo summarizing flanking-base composition.

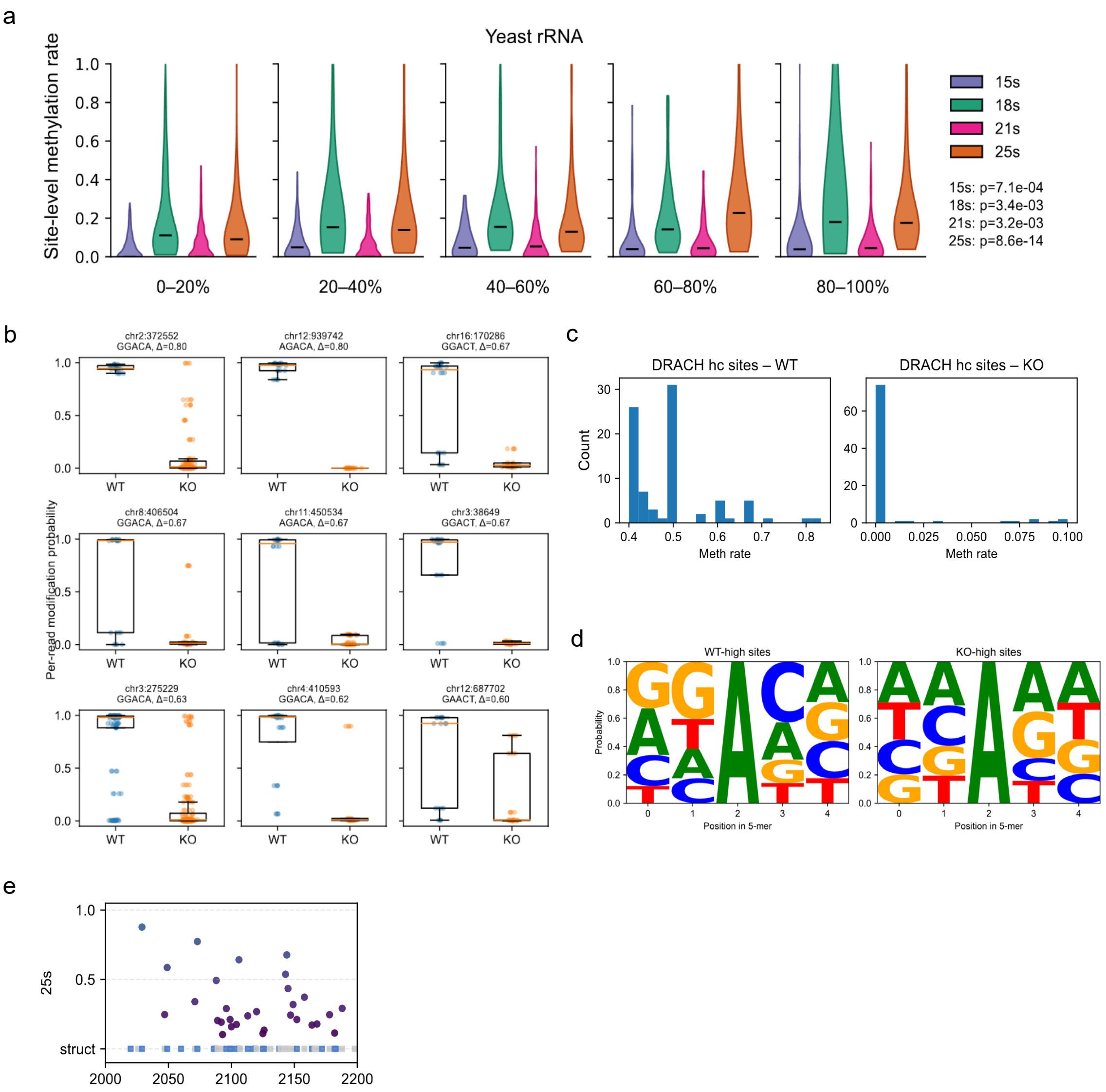


**Figure S17:** Positional and secondary-structure context of m$^6$A sites in *S. cerevisiae* rRNA. **a**, Distributions of site-level m$^6$A rates along yeast rRNAs, summarized across relative position bins for each rRNA species. **b**, Comparison of m$^6$A rates at representative DRACH sites between wild-type (WT) and methyltransferase-deficient knockout (KO) strains across replicates. **c**, Global distributions of site-level m$^6$A rates for DRACH sites in WT and KO, highlighting shifts in modification levels. **d**, Sequence logos of flanking 5-nt contexts for sites enriched in WT and KO, summarizing context differences associated with condition-specific modification. **e**, Local view of a representative rRNA segment showing secondary-structure categories together with site-level m$^6$A rates, illustrating the relationship between modification and structural context.

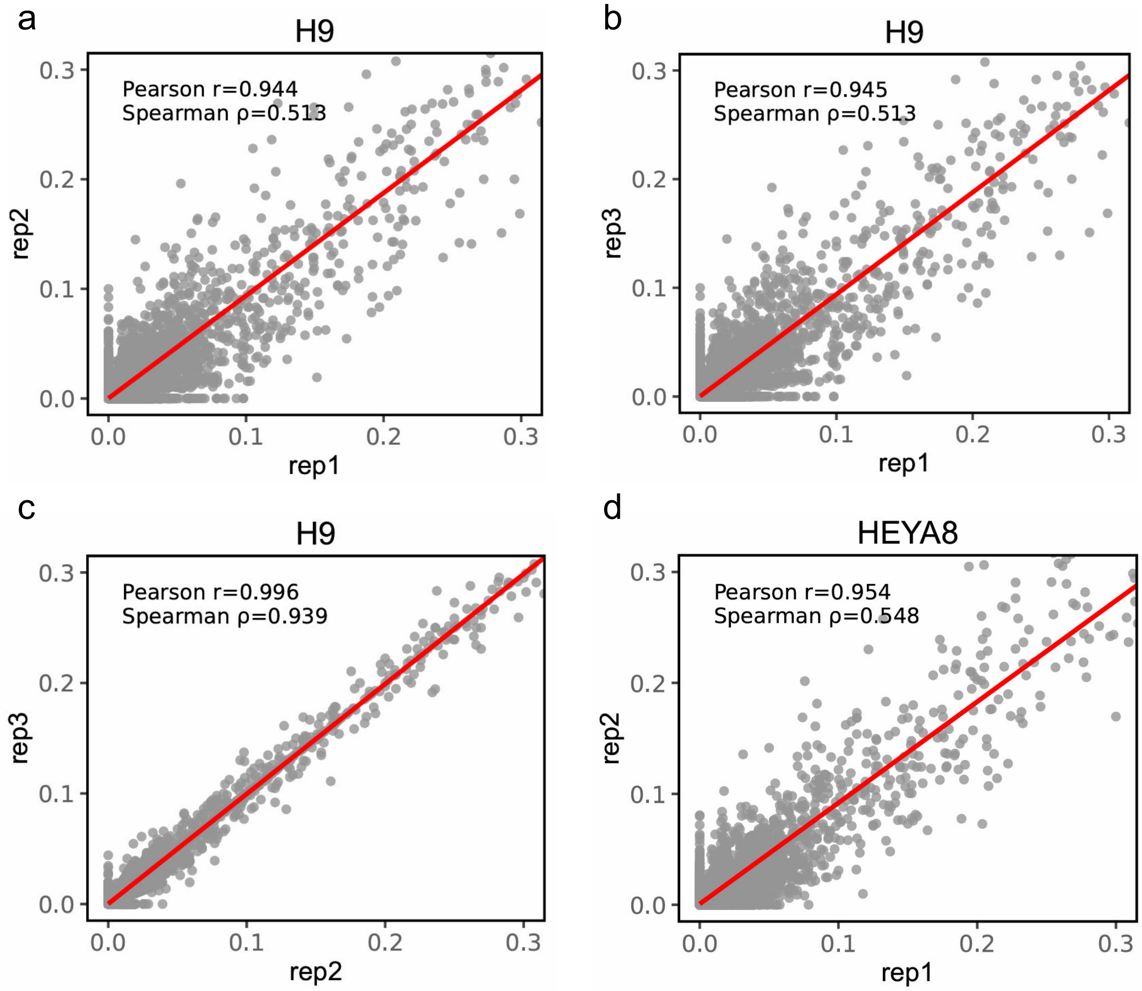


**Figure S18:** Cross-batch reproducibility of predicted modification levels in H9 and HEYA8 cell lines. **a**, Scatter comparison of predicted site-level modification levels between H9 replicate 1 (rep1) and replicate 2 (rep2), with Pearson $r$ and Spearman $\rho$ indicated. **b**, Scatter comparison between H9 rep1 and rep3. **c**, Scatter comparison between H9 rep2 and rep3. **d**, Scatter comparison between HEYA8 rep1 and rep2. Each point represents the predicted modification level of the same site across two replicates, and the red diagonal line indicates $y = x$.

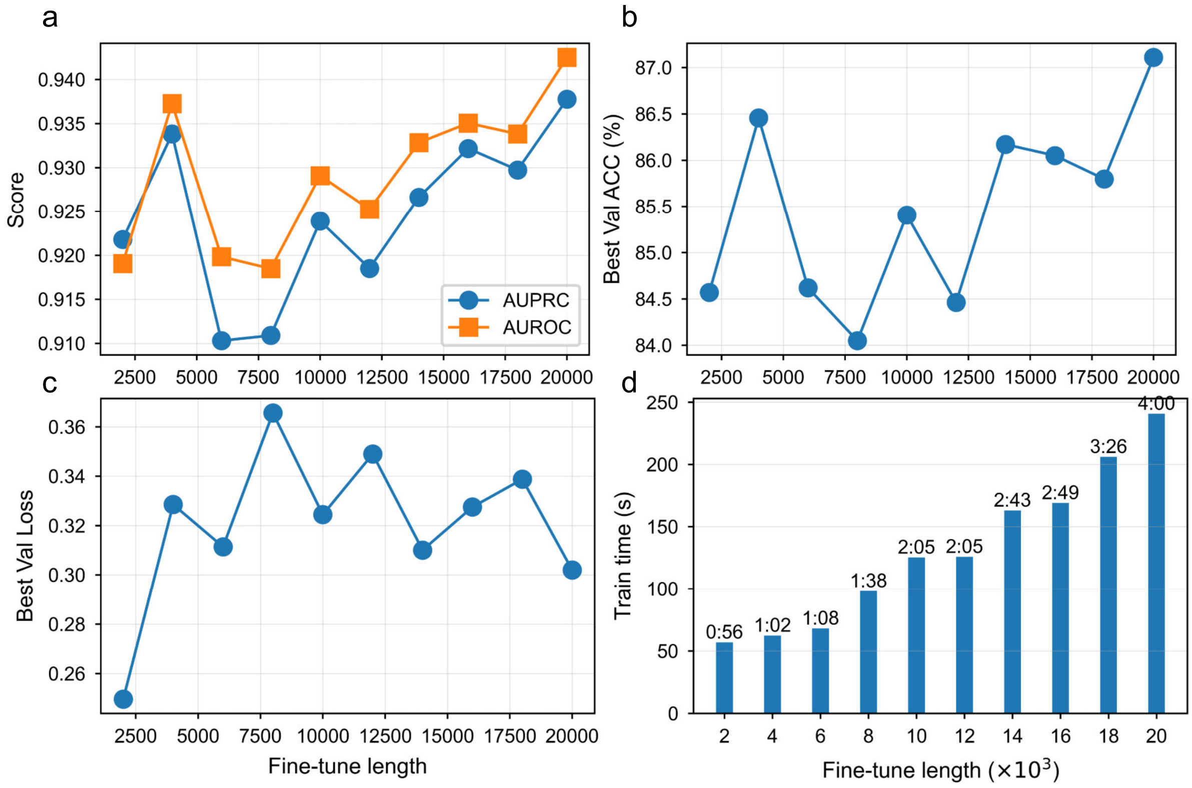


**Figure S19:** Impact of fine-tuning set size on $m^5U$ model performance and training cost. **a**, AUPRC and AUROC across different fine-tuning set sizes. **b**, Best validation accuracy across different fine-tuning set sizes. **c**, Best validation loss across different fine-tuning set sizes. **d**, Training time across different fine-tuning set sizes, with time labels shown above each bar in mm:ss format.

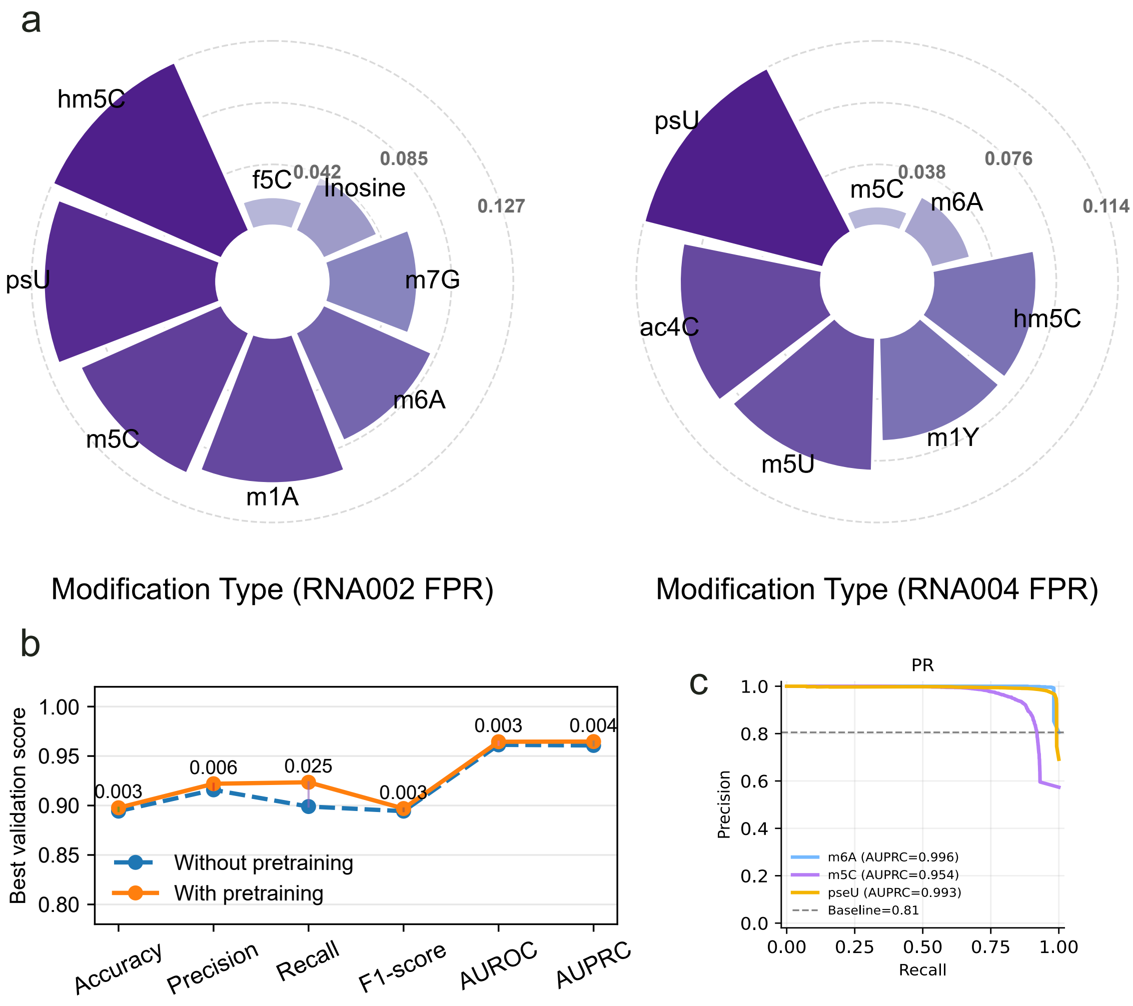


**Figure S20:** False positive rate profiles and pretraining-associated performance gains. **a**, Circular bar plots showing false positive rates (FPRs) across RNA modification types under RNA002 and RNA004 settings. The radial axis indicates the FPR value for each modification type. **b**, Best validation scores of models trained with or without pretraining across accuracy, precision, recall, F1-score, AUROC and AUPRC. Numeric labels denote the absolute improvement obtained after pretraining.**c**, Precision–recall curves of Dorado RNA004 modified-base calling for $m^6A$, $m^5C$ and pseudouridine (pseU), using the corresponding modified samples as positives and the unmodified sample as the negative control.

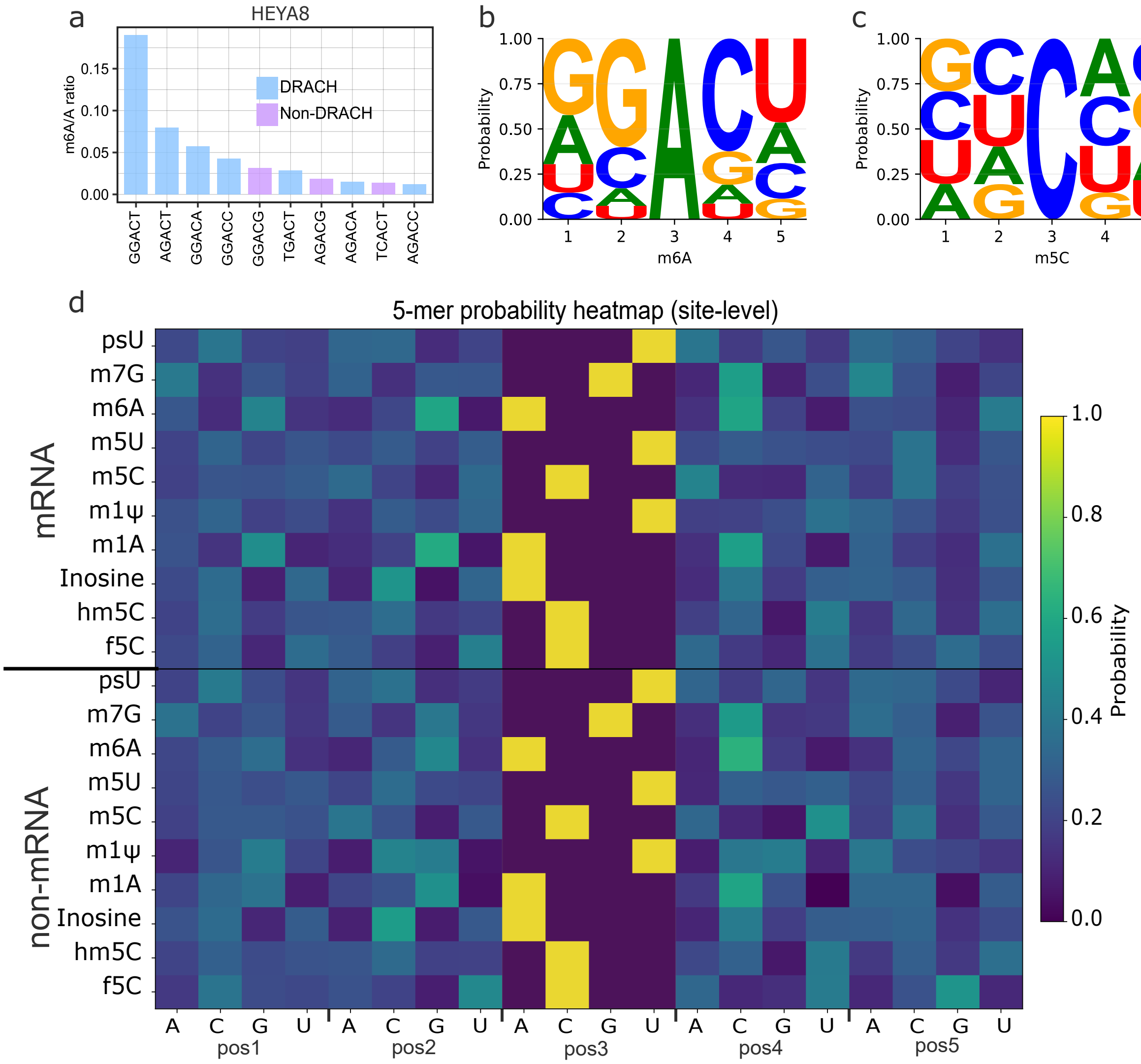


**Figure S21:** Motif preference of m$^6$A in cells and sequence-context signatures across ten RNA modifications. **a**, Enrichment of m$^6$A calls across representative 5-mers in HEYA8, stratified by canonical DRACH status (DRACH versus non-DRACH). **b**, Sequence logo of the local 5-nt context (±2 nt; A-centered 5-mer) surrounding m$^6$A sites, highlighting nucleotide preferences around modified adenosines. **c**, Sequence logo of the local 5-nt context (±2 nt; C-centered 5-mer). **d**, Site-level 5-mer probability heatmap summarizing base preferences for ten RNA modifications, shown separately for mRNA and non-mRNA regions. Columns indicate 5-mer position (pos1–pos5) and nucleotide (A/C/G/U), and colours represent the corresponding probability (0–1).

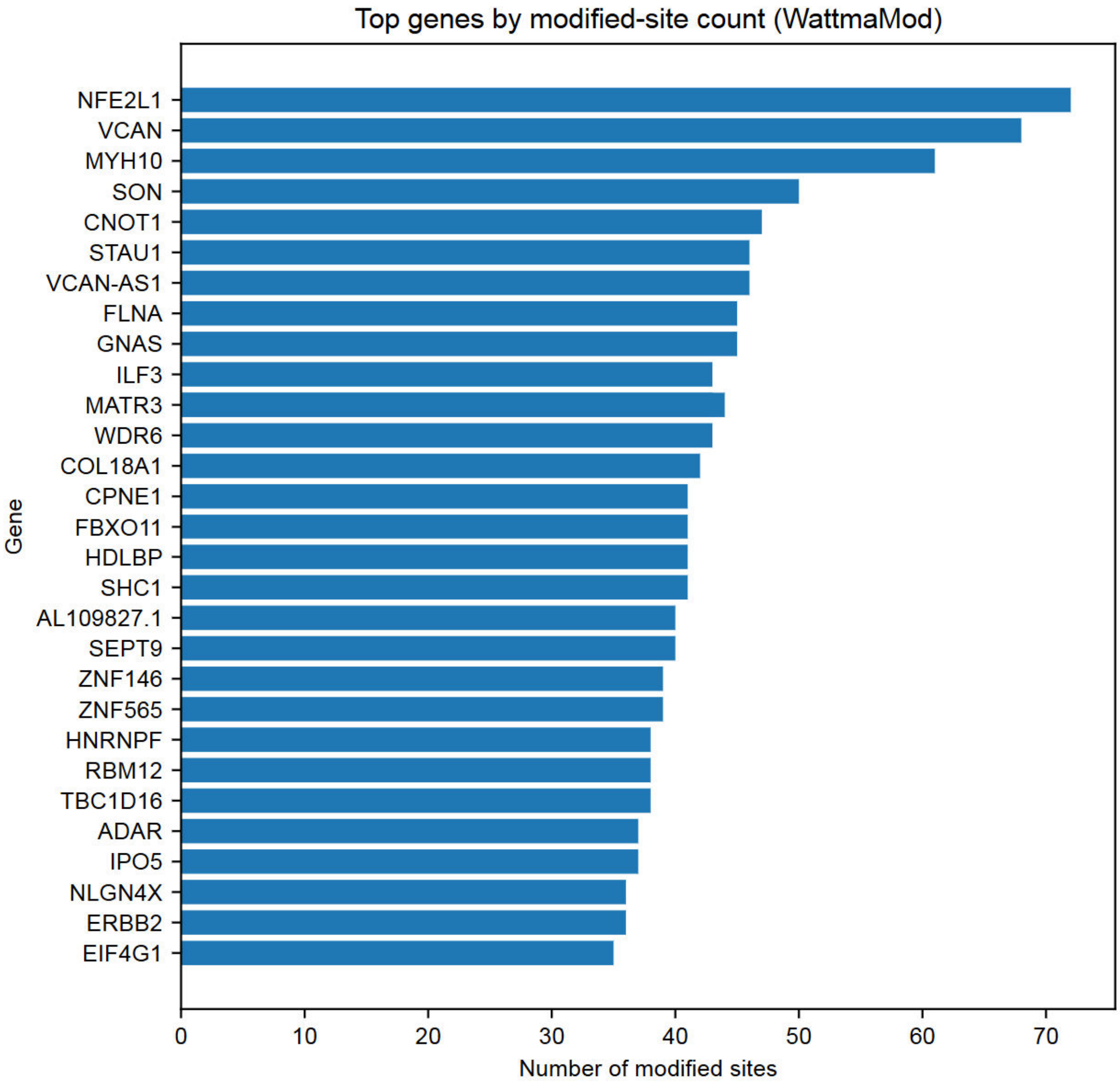


**Figure S22:** Gene-level distribution of WattmaMod-predicted $m^6A$ sites detected under the RRACH motif in H9 cells.

Bar plot showing the top 30 genes with the highest numbers of predicted modified sites under the RRACH motif.

### Supplementary Training Configuration

#### Model and training setup

This framework comprises wavelet-guided[1] multiscale convolutions, channel-level attention mechanisms[2], event feature sequence modeling, and state space sequence encoders[3]. The signal undergoes discrete wavelet multiscale decomposition, with low-frequency components resampled via linear interpolation and high-frequency components via nearest-neighbor interpolation. These components then pass through a dual one-dimensional convolutional layer configured as (5, 64, 64) and a channel attention module.

Event feature branches are encoded via a bidirectional LSTM[4] with hidden size 64. Subsequently, signals are fused with events through modeling using a dynamic gated cross-attention mechanism and Mamba's sequence module. Projection is performed via a multi-layer feedforward projection head with dimensions 768,512,256,256. The final classifier consists of two layers of multilayer perceptron heads: Linear(256,128) and Linear(128,num_classes=2).

[3] Albert Gu and Tri Dao. Mamba: Linear-time sequence modeling with selective state spaces. In *First conference on language modeling*, 2024.

[4] Alex Graves and Jürgen Schmidhuber. Framewise phoneme classification with bidirectional lstm and other neural network architectures. *Neural networks*, 18(5-6):602–610, 2005.

**Table S3:** Hyperparameter settings and training configuration.

| **Module** | **Parameter** | **Argument** | **Value** |
|---|---|---|---|
| Self-supervised augmentation | Jitter ratio | `jitter_ratio` | 0.1 |
| | Scaling ratio | `scaling_ratio` | 0.1 |
| | Max masked segments | `max_segments` | 5 |
| | Frequency mask ratio | `frequency_mask_ratio` | 0.1 |
| | Frequency add ratio | `frequency_add_ratio` | 0.1 |
| Supervised augmentation (signal) | Time-warp sigma | `time_warp_sigma` | 0.005 |
| | Time-warp knots | `time_warp_knots` | 2 |
| | Magnitude-warp sigma | `mag_warp_sigma` | 0.005 |
| | Magnitude-warp knots | `mag_warp_knots` | 2 |
| | Number of augmented views | `n_views` | 6 |
| Supervised contrastive | Margin | `margin` | 0.5 |
| | Temperature | `temperature` | 0.1 |
| | Projection dim | `proj_dim` | 128 |
| Model (WattmaMod) | Model width | `d_model` | 64 |
| | State size | `d_state` | 16 |
| | Convolution width | `d_conv` | 2 |
| | Dropout | `dropout` | 0.3 |
| Input | k-mer length | `k` | 5 |
| Signal preprocessing | Interpolation length | $L_{\text{interp}}$ / `interp_len` | 500 |
| Optimization | Optimizer | `optimizer` | AdamW |
| | Initial learning rate | `lr` | 0.001 |
| | Weight decay | `weight_decay` | 1e-5 |
| | Batch size | `batch_size` | 512 |
| | Epochs (pretrain) | `epochs_pretrain` | 50 |
| | Epochs (finetune) | `epochs_finetune` | 20 |
| Scheduler (ReduceLROnPlateau) | Mode | `mode` | `min` |
| | Factor | `factor` | 0.5 |
| | Patience (epochs) | `patience` | 3 |
| | Threshold | `threshold` | $1 \times 10^{-6}$ |
| | Minimum LR | `min_lr` | $1 \times 10^{-6}$ |
| Training stabilization | Gradient clipping norm | `max_norm` | 1.0 |
